# Single cell quantification of ribosome occupancy in early mouse development

**DOI:** 10.1101/2021.12.07.471408

**Authors:** Tori Tonn, Hakan Ozadam, Crystal Han, Alia Segura, Duc Tran, David Catoe, Marc Salit, Can Cenik

**Author notes:** These authors contributed equally (Tori Tonn, Hakan Ozadam, Crystal Han).

## Abstract

Technological limitations precluded transcriptome-wide analyses of translation at single cell resolution. To solve this challenge, we developed a novel microfluidic isotachophoresis approach, named RIBOsome profiling via IsoTachoPhoresis (Ribo-ITP), and characterized translation in single oocytes and embryos during early mouse development. We identified differential translation efficiency as a key regulatory mechanism of genes involved in centrosome organization and N^6^-methyladenosine modification of RNAs. Our high coverage measurements enabled the first analysis of allele-specific ribosome engagement in early development and led to the discovery of stage-specific differential engagement of zygotic RNAs with ribosomes. Finally, by integrating our measurements with proteomics data, we discovered that ribosome occupancy in germinal vesicle stage oocytes is the predominant determinant of protein abundance in the zygote. Taken together, these findings resolve the long-standing paradox of low correlation between RNA expression and protein abundance in early embryonic development. The novel Ribo-ITP approach will enable numerous applications by providing high coverage and high resolution ribosome occupancy measurements from ultra-low input samples including single cells.

## INTRODUCTION

Temporal regulation of gene expression is critical for embryonic development. The early gene expression landscape is shaped by post-transcriptional regulation of maternal transcripts due to the absence of transcription from later stages of oocyte maturation through the early divisions of the embryo^1^. Consequently, RNA expression and protein abundance are only modestly correlated until the late morula stage emphasizing the need to elucidate post-transcriptional regulation during initial stages of embryogenesis^2–4^. In particular, translational control of specific transcripts is essential for oocyte maturation and the oocyte-to-embryo transition^5, 6^. Yet, the transcriptome-wide characterization of translation in early mammalian embryonic development has remained unexplored to date despite the existence of ribosome profiling techniques for more than a decade^7^.

Ribosome profiling provides a snapshot of transcriptome-wide mRNA translation by high-throughput sequencing of RNA fragments protected by ribosomes from nuclease digestion^8^. However, the conventional ribosome profiling approach involves multiple steps with significant loss of input material restricting its application to samples with large numbers of cells. Consequently, many important questions related to translational control remain to be addressed due to limited availability of biological material.

To overcome this constraint, we developed a novel method leveraging the principles of microfluidic on-chip isotachophoresis (ITP) for isolation of ribosome protected fragments (RPFs). ITP has previously been applied for extraction of nucleic acids from blood, urine, and cell culture samples^9^. Compared to conventional RNA extraction approaches, ITP offers faster processing times, no requirement of liquid transfers, and high yield with low RNA inputs^10, 11^. Despite these advantages, ITP is considered to lack the ability to deliver the stringent size selection that would be required for applications such as ribosome profiling^12, 13^.

## Results

### RIBOsome profiling via IsoTachoPhoresis (Ribo-ITP): an on-chip method for high yield extraction and efficient library preparation of ribosome protected fragments

Here, we designed and manufactured a custom microfluidic polydimethylsiloxane (PDMS) chip to recover ribosome footprints from nuclease-digested lysates with high yield using a novel, specialized technique named RIBOsome profiling via IsoTachoPhoresis (Ribo-ITP) (Fig. 1A, 1B, Extended Data Fig S1; Supplementary Video 1). We implemented numerous innovations that enable a chemistry required to achieve single-cell ribosome profiling by coupling ITP with an optimized on-chip size selection. Specifically, we leveraged pretreatment of the channel with benzophenone to enable light-induced polymerization of polyacrylamide inside PDMS chips^14^. To aid visualization, we included DNA oligonucleotide markers containing a 5’ fluorophore and 3’ dideoxycytosine modification to prevent marker amplification in downstream library preparation (Extended Data Fig. 2). An on-chip buffer exchange allowed the purified RNAs to be directly compatible with 3’ dephosphorylation, the first step in sequencing library preparation (Extended Data Fig. 3). Finally, we adopted an efficient single tube library preparation chemistry that relies on a template switching reverse transcriptase and incorporation of unique molecular indexes (UMIs) at the 5’ end of the RPFs. Collectively, these optimizations reduce sample requirements by many orders of magnitude while simultaneously reducing sample processing time to deliver ribosome occupancy measurements from ultra-low input samples, including single cells.

**Figure 1.**
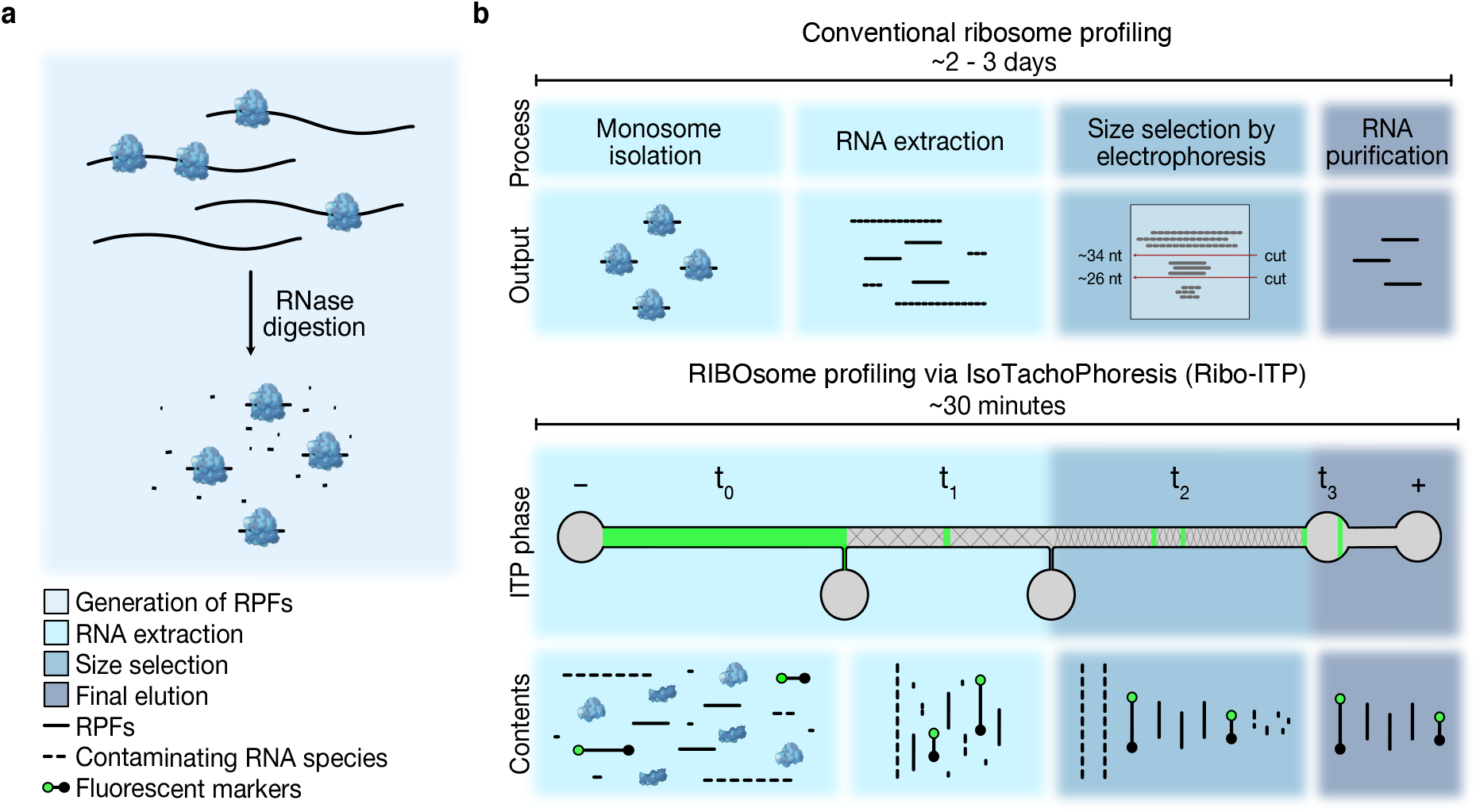
Schematic of Ribo-ITP. **a,** Schematic of the generation of ribosome protected fragments (RPFs). Following RNase digestion, RPFs are isolated with the conventional or novel Ribo-ITP approach. **b,** Schematic of the conventional ribosome profiling protocol and the Ribo-ITP process for extraction of RPFs. In Ribo-ITP, marker oligonucleotides with a 5’ fluorophore and 3’ ddC blocking modification, which encapsulate the size range of RPFs, are added to the digested cellular lysate. Lysate contents are loaded into the channel (t0), then an electrical current is applied to selectively focus species of a specific electrophoretic mobility range, enabling nucleic acid extraction by isotachophoresis. Nucleic acids are extracted in a narrow ITP band, and then size selected as they migrate through 5% (t1) and 10% (t2) polyacrylamide gels, respectively. At the end of the run, purified and size-selected RNAs are collected (t3).

### Ribo-ITP enables simultaneous extraction, stringent size selection and high yield recovery of RNAs from ultra-low input samples

In the conventional ribosome profiling method, RPFs are isolated by phenol-chloroform based RNA extraction followed by size selection using polyacrylamide gel electrophoresis^15^. Given that a typical mammalian cell contains ∼10-40 pg of RNA, an approach capable of generating ribosome occupancy measurements from such limiting amounts needs to maintain consistently high yield of RPF recovery with inputs in the picogram range.

We first compared the recovery of RNAs that span the typical size range of RPFs (∼21-35 nt) achieved by the conventional method^16^ versus our new approach, Ribo-ITP. When using 20 ng input samples, Ribo-ITP yielded 94 ± 3.5% (SEM) recovery in contrast to only 38 ± 10.9% (SEM) for the conventional gel extraction approach (Extended Data Fig. 4). We then adopted a radioactive labeling assay to visualize and quantify the recoveries from ultra-low RNA inputs (40 pg - 2 ng). With a 2 ng RNA input, 87.5 ± 3.2% yield was achieved by Ribo-ITP compared to 35.3 ± 11.4% by conventional gel extraction (Fig. 2A). When RNA inputs were decreased further to 400 pg and 40 pg, the recovery by Ribo-ITP remained high at 74 ± 6.1% and 67.5 ± 10.6%, respectively (Fig. 2A). Gel extraction had negligible yield with these samples. Thus, the consistently high RNA yields obtained with Ribo-ITP demonstrate that this method empowers high yield extraction even at ultra-low inputs.

**Figure 2.**
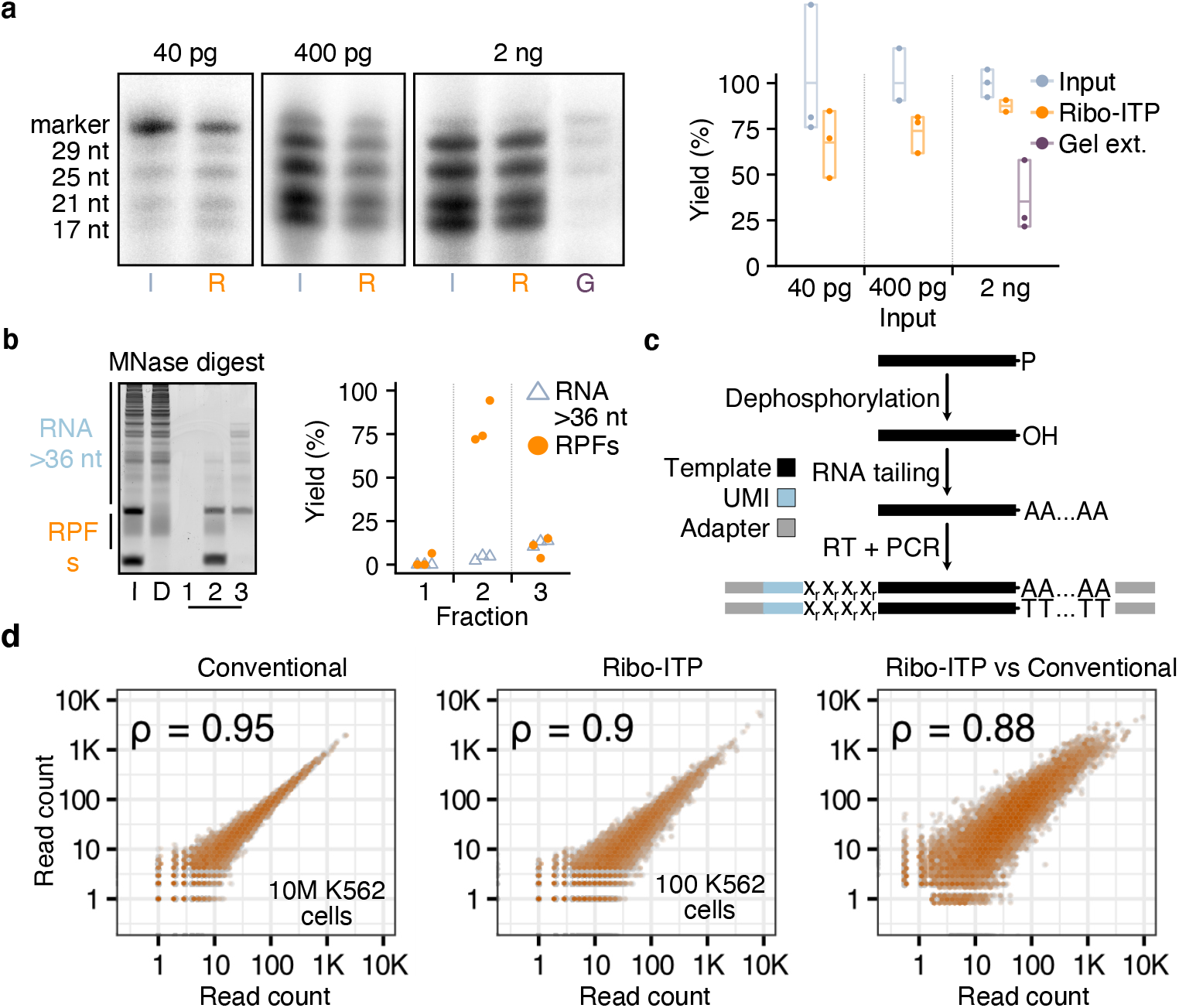
Characterization of Ribo-ITP method and validation of efficacy in ultra-low input ribosome profiling. **a,** Representative gel images highlighting inputs (I), RNAs recovered by Ribo-ITP (R), and gel electrophoresis (G) are shown. Four RNAs of 17, 21, 25, and 29 nt used in the experiment were radioactively labeled at their 5’end. Percent yield was calculated for the 25 nt RNA. **b,** Representative gel image of a size selection experiment. 100 ng of MNase-digested RNA from K562 cells (D) was used as an input for Ribo-ITP after the addition of the two fluorescent marker oligonucleotides (I). In a typical experiment, we collect the sample flanked by the two fluorescent nucleotide markers (Fraction 2 here). Here, we also collected the RNAs that eluted before the arrival of the shorter fluorescent marker (Fraction 1) as well as the RNAs that were located behind the longer fluorescent marker (Fraction 3), which typically remain in the channel. The percent yield of RNAs larger than the longer fluorescent marker oligonucleotide (> ∼36 nt) (blue) and RNAs flanked by the markers (orange), corresponding to the size range of RPFs, are plotted for each fraction. **c,** Schematic of the sequencing library preparation protocol. In a single-tube reaction, isolated RPFs are 3’ dephosphorylated and poly(A)-tailed. A template-switching reverse transcriptase (RT) creates templates that incorporate UMI-containing adapters. **d,** Pairwise correlation of gene-level ribosome occupancy measured in conventional ribosome profiling and Ribo-ITP from human K562 cells. The left plot highlights two replicates of conventional ribosome profiling experiments from ∼10M cells. The middle plot is from two replicates of Ribo-ITP with ∼100 cells. For the plot on the right, we used the mean number of counts per million reads for each gene. The Spearman correlation coefficients between the gene-level ribosome occupancies are indicated on the top left corner.

To analyze the efficiency of our method at excluding RNA fragments larger than RPFs (>36 nt), we digested total RNA from a human myelogenous leukemia cell line (K562) with micrococcal nuclease (MNase), purified the sample, and subjected it to Ribo-ITP (Methods; Fig. 2B). We achieved 94% exclusion of the unwanted large RNA fragments (>36 nt) (Fig. 2B). Finally, to verify the ability of Ribo-ITP to extract RNAs from complex cellular lysates, we spiked RPF-sized synthetic RNAs (17, 21, 25, 29 nt) into total cellular lysates from ∼1000 K562 cells. Ribo-ITP of this sample recovered the spiked RNAs with stringent size selection and high yield (Extended Data Fig. 5). Collectively, these results indicate that Ribo-ITP can simultaneously extract and size-select RPF-size RNAs from cellular lysates with high yield.

### Ribo-ITP enables high quality single cell ribosome occupancy measurements

A critical factor in generating ribosome profiling data is the choice of the ribonuclease^17^. Here, we employed micrococcal nuclease (MNase) to maximize the recovery of ribosome protected RNA fragments^18–20^ at the expense of a slight loss in nucleotide resolution, which can be ameliorated by computational means^21^. To validate the quality of ribosome profiling data, we first performed Ribo-ITP from 100 K562 cells and conventional ribosome profiling using the gold-standard method of monosome isolation^15^ from 10 million K562 cells. For sequencing library preparation, we optimized a single tube protocol that incorporates unique molecular identifiers (UMIs) via a template switching reverse transcriptase (Fig. 2C).

Ribosome occupancy measurements from 100 cells obtained using Ribo-ITP were highly reproducible across replicates (Fig. 2D; Extended Data Fig. 6; Supplementary Table 1). The footprints displayed the characteristic read length distribution^22^ and the expected enrichments at annotated translation start and stop sites (Extended Data Fig. 7). The vast majority of transcript mapping reads originated from the coding regions and were highly enriched over the distribution expected from random fragmentation (Chi-squared test, p-value < 2.2 x 10^-16^; Extended Data Fig. 8). Critically, ribosome profiling measurements from 100 cells generated by Ribo-ITP recapitulated the conventional ribosome profiling measurement (Spearman correlation coefficient 0.88; p-value < 2.2 x 10^-16^, Fig 2D). These results reveal that ribosome occupancy can be accurately measured from as few as 100 human cells using Ribo-ITP.

Next, we applied Ribo-ITP and RNA-seq to characterize the translation changes of single cell and single embryos in the context of early embryonic development in mice. In particular, the initial division of zygotes occurs in the absence of new RNA synthesis, rendering the translation of stored maternal transcripts absolutely essential for the early stages of development. Hence, we generated Ribo-ITP and RNA-seq across multiple stages of preimplantation development including single unfertilized oocytes at germinal vesicle (GV) and metaphase II (MII) stages, as well as single fertilized embryos from the 1-cell zygote to 8-cell stages (Fig. 3A; Supplementary Table 1).

**Figure 3.**
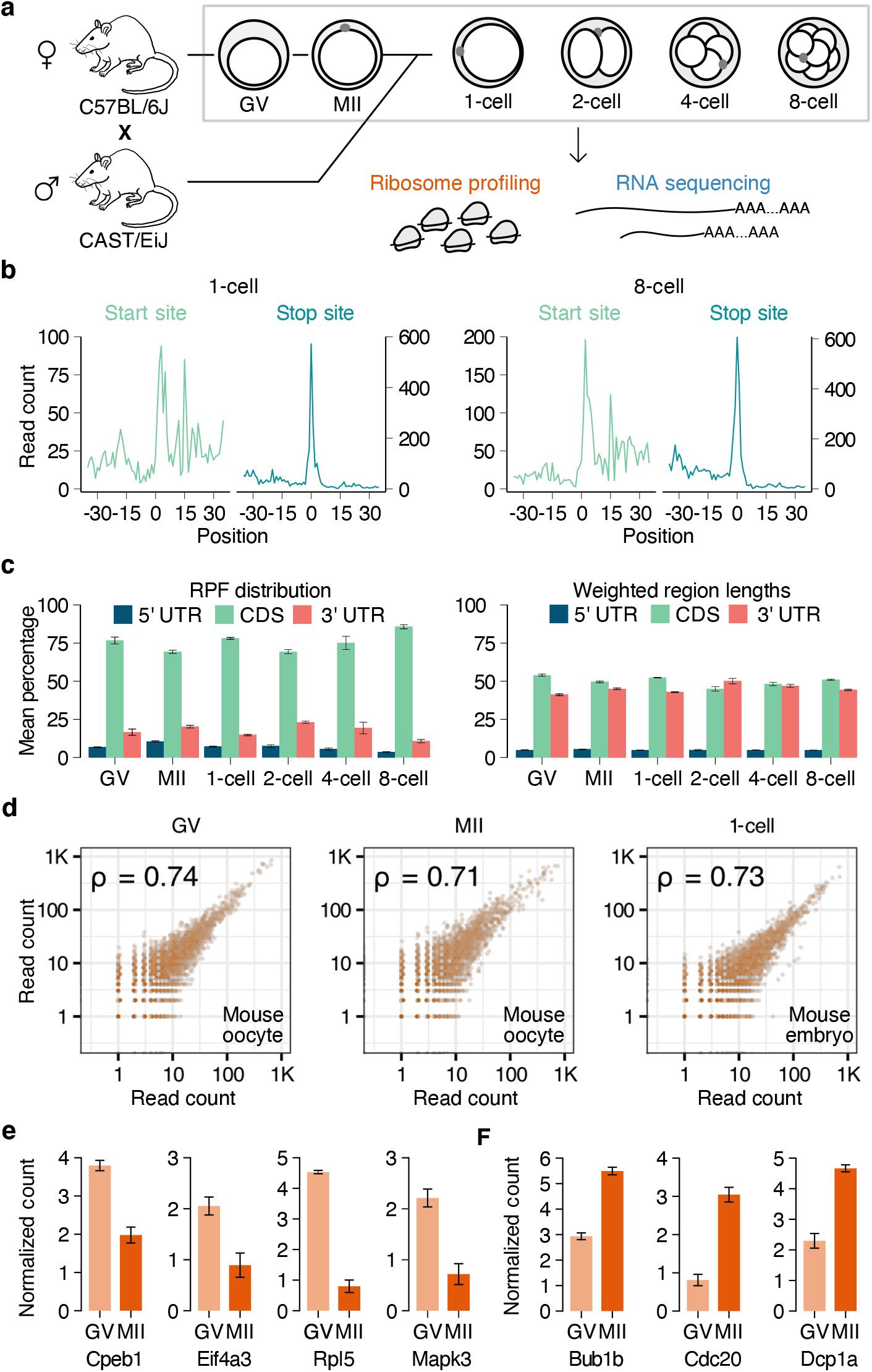
Ribo-ITP enables single cell and single embryo measurements of ribosome occupancy. **a,** Schematic of the mouse experiments. Unfertilized oocytes (GV- and MII-stage) from C57BL/6J strain along with zygotes to the 8-cell stage embryos from a crossbreed of two strains (C57BL/6J and CAST/EiJ) were collected for RNA expression and ribosome occupancy measurements. **b,** Ribosome occupancy around the translation start and stop sites in a representative zygote (1-cell) and an 8-cell stage embryo. Translation start (or stop) sites are denoted by the position 0. Aggregated read counts (y-axis) relative to the start (or stop) sites are plotted after A-site correction (Methods). **c,** On the left, the distribution of reads across transcript regions (5’ UTR, CDS and 3’UTR) are shown. On the right, the distribution of the lengths of these regions weighted by their ribosome occupancy are depicted. The error bars indicate the standard error of the mean percentages. **d,** Pairwise correlation of gene-level ribosome occupancy in single cells are plotted along with Spearman correlation coefficients (top left). **e,f,** The standard error and mean of centered log ratio of the ribosome occupancy (y-axis) was plotted for representative transcripts that were previously shown to have increased polysome association in GV- (panel E) or MII-stage (panel F) oocytes23 (remaining genes are shown in Fig. S12).

In the single cell ribosome occupancy data, we observed a median of 48,017 unique molecules originating from the coding regions of transcripts, leading to detection of an average of 5064 genes per cell (range 4076 - 6679; Extended Data Fig. 9). Furthermore, single oocyte and single embryo ribosome profiling data demonstrated the expected enrichment of footprints mapping to coding regions and characteristic enrichments at the start and stop sites (Fig. 3B, 3C, Extended Data Fig. 10). Replicate measurements of ribosome occupancy had high correlation with each other (Fig. 3D; Extended Data Fig. 11).

To validate the quality of our single cell ribosome profiling measurements, we compared our results to a complementary approach for assaying translation; polysome profiling, a strategy that can inform the distribution of the number of ribosomes on a given mRNA. In particular, a previous study collected ∼500-600 GV- and MII-stage oocytes and validated changes in polysome association with rt-qPCR experiments for 29 transcripts^23^. Remarkably, our single cell ribosome profiling measurements recapitulated the previously identified changes in polysome association for 28 out of 29 RNAs (Fig. 3E, Extended Data Fig. 12; Methods). Taken together, our results indicate that Ribo-ITP enables highly consistent and high quality ribosome occupancy measurements from single cells and single embryos during early mouse development.

### Ribo-ITP reveals allele-specific translation in mouse preimplantation development

In mouse development, transcription of the zygotic genome is activated at the 2-cell stage. Yet, we currently do not know when newly synthesized RNAs engage with ribosomes and whether there exist any gene- and allele-specific differences in these dynamics. We first addressed the question of allele-specific expression following zygotic genome activation. Both deterministic and stochastic differences in allele expression ratios are believed to contribute to differentiation and normal development, though studies had been limited to the level of epigenetics and transcription in the early mouse embryo^24, 25^. Since these studies typically require single-cell resolution, it has been impossible to study allele-specific translation until now.

To distinguish RNAs derived from the maternal and paternal alleles, we analyzed embryos from a cross of two mouse strains (C57BL/6J × CAST/EiJ). Using strain-specific single-nucleotide polymorphisms (SNPs) to distinguish maternal and paternal RNAs, we detected 229,991 unique parent-of-origin-specific RPFs mapping to coding regions (Methods). As a control for the accuracy in alignments and SNP annotation, we analyzed unfertilized MII-stage oocytes and found 97.3% correctly classified reads, i.e., reads with maternal SNPs, in our ribosome profiling experiments.

To monitor allele-specific ribosome engagement alongside corresponding RNA expression^26^, we specifically focused on the paternal allele which, unlike RNA of maternal origin, is a proxy of newly synthesized transcripts (Methods). We analyzed the global pattern of ribosome engagement of paternally-derived RNAs, i.e., paternal allele ratios, by aggregating reads across all detected genes. We found that, coinciding with the activation of zygotic transcription, the paternal ratio of ribosome occupancy steadily increased from 7.1% in the 2-cell stage to 47.7% in the 8-cell stage embryos (Fig. 4A; Extended Data Fig. 13). Importantly, we discovered that the ratio of paternal alleles across these stages was statistically indistinguishable between ribosome occupancy and RNA expression (t-test; p-value > 0.14 for all stages; Fig. 4A). This result indicates that ribosome engagement is overall concurrent with the synthesis of paternal RNAs via zygotic genome activation.

**Figure 4.**
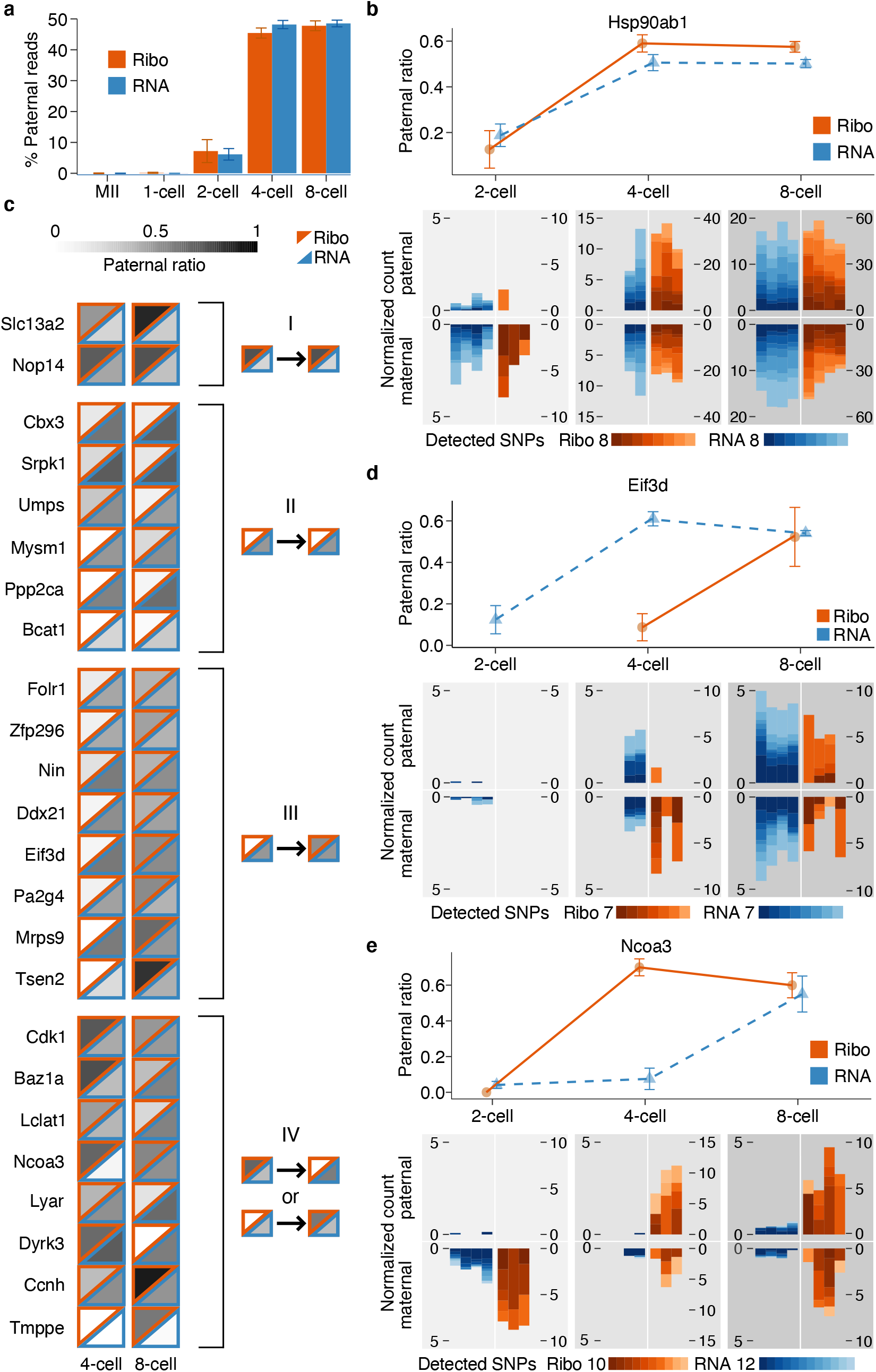
Allele specific translation and RNA expression in early mouse development. **a,** Strain-specific SNPs were used to assign sequencing reads to the paternal and maternal allele (Methods). Standard error and mean of the percentage of paternal reads in each stage (y-axis) is plotted. **b,d,e,** Line-plots (top) indicate the percentage of paternal reads (y-axis) in RNA-Seq and Ribo-ITP experiments. The reads are combined across replicates and error bars indicate standard error of the mean of paternal ratios. At the bottom, maternal and paternal reads counts per 10k are plotted for all individual replicates and SNPs. The total number of detected coding SNPs and their corresponding colors are shown with color scales. **c,** The percentage of paternal reads is indicated by different shades of gray for ribosome occupancy (upper triangles with orange borders) and RNA expression (lower triangles with blue borders). Genes that displayed differential ribosome engagement in an allele-specific manner in comparison to RNA expression were grouped into four clusters. The prototypical shared pattern in each cluster is displayed on the right.

We next considered whether there are any gene-specific exceptions to the observed global pattern of equal paternal allelic ratios in RNA expression and ribosome occupancy in the early mouse embryo. Our high coverage data enabled us to assess these dynamics in 1012 genes (Methods). As expected, the majority of genes exhibited a similar ratio of paternal reads in both RNA expression and ribosome occupancy (Extended Data Fig. 14). For example, we detected 8 distinct coding SNPs differentiating the two alleles in *Hsp90ab1* across multiple replicates in RNA-seq and Ribo-ITP. The high similarity of paternal allele ratio in RNA expression and ribosome occupancy was consistently observed for multiple replicates and supported by distinct SNPs, highlighting the reproducibility and high coverage of our allele-specific measurements (Fig. 4B, Extended Data Fig. 15).

We identified 24 genes that had differential ribosome engagement in an allele-specific manner in comparison to RNA expression (two-sample test for the equality of proportions; see Methods for details; Fig. 4B). We identified four clusters among these 24 genes (Fig. 4C). While clusters I and II encompass genes that display consistent allele-specific ribosome occupancy bias throughout early development (Extended Data Fig. 16A), genes in the other two clusters display allele-specific ribosome occupancy in a stage-dependent manner.

In particular, several genes including *Eif3d* display delayed engagement of newly transcribed paternal RNA with ribosomes. Specifically, paternal allele was robustly expressed in 4-cell embryos, yet ribosome association of the paternal allele is delayed until the 8-cell stage (Fig. 4E; Cluster III, Extended Data Figs. 16B-C). This observation suggests that specific transcripts may either have slow kinetics of nuclear export, or are sequestered in translationally inactive compartments until their subsequent association with ribosomes occurs in the 8-cell stage.

Genes in the last group (Cluster IV) included *Cdk1*, a key regulator of cell cycle, *Baz1a*, a chromatin remodeling factor, and *Lclat1*, lysocardiolipin acyltransferase 1 (Fig. 4D; Extended Data Figs. 16D-F). While some of these genes had differential ribosome occupancy of the paternal allele compared to RNA expression in only the 4-cell stage (e.g. *Baz1a*), others (e.g. *Lclat1*) differed at the 8-cell stage. These findings suggest the presence of an interaction between the cis-elements and regulatory factors that enable differential ribosome occupancy of one of the alleles only in a specific stage of development. Taken together, our results reveal that for most transcripts, ribosome engagement is concurrent with zygotic activation and paternal RNA expression. Yet, a small number exhibit allele-specific ribosome engagement including stage-specific differences.

### Differential translation efficiency during oocyte maturation and preimplantation development regulates a functionally coherent set of genes

We next characterized transcript-specific changes in translation across the studied developmental stages. Transcripts with the highest variability of ribosome occupancy revealed two major transitions: one between GV- and MII-stage oocytes and another between 2-cell and 4-cell stage embryos (Fig. 5A; see Methods). We then focused on identifying the set of transcripts with differences at the translational level, i.e., with differential translation efficiency, as defined by significant changes in ribosome occupancy while controlling for RNA abundance (Methods).

**Figure 5.**
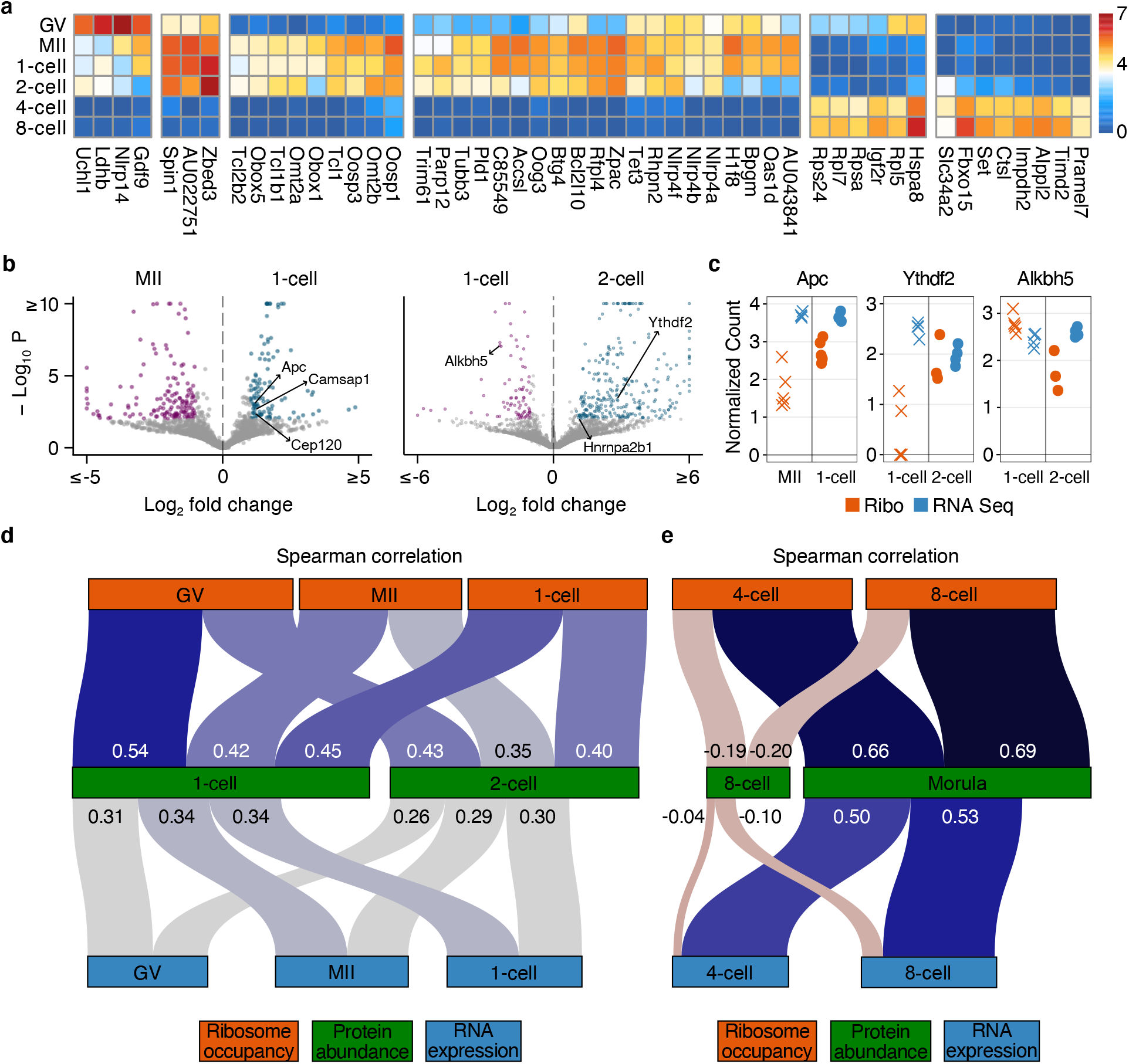
Differential translation efficiency between developmental stages and association between ribosome occupancy and protein abundance. **a**, 50 genes with the highest variability in ribosome occupancy across developmental stages are plotted (Methods). The colors correspond to the average of the centered log-ratio of ribosome occupancy where average is taken across replicates. **b**, Volcano plots depict the statistical significance (y-axis) and log2 fold-change (x-axis) in translation efficiency between two developmental stages (Methods). Colored points indicate transcripts with significant differences (FDR <0.01). **c**, The centered log-ratio normalized read counts from Ribo-ITP and RNA-seq experiments are plotted for the highlighted genes. All replicate measurements from the given developmental stage are shown. **d, e**, Sankey diagrams depict the relationships between protein abundance with RNA expression and ribosome occupancy. The color and thickness of the links connecting the nodes are proportional to the strength of the Spearman rank correlation (Methods).

We uncovered a large number of genes that exhibited translation efficiency changes during oocyte maturation from GV- to MII-stage, as well as upon fertilization (Fig. 5B; Extended Data Fig. 17; Supplementary Table 2). Strikingly, among the 129 genes that were translationally upregulated upon fertilization, 126 had no statistically significant changes in RNA expression (FDR 0.01; 1-cell embryo vs MII-stage oocyte; Fig. 5B; 5C). These genes were significantly enriched for cytoskeleton organization (Fisher’s exact test odds ratio 4.67; p-value = 8.5 x 10^- 09^; Supplementary Table 3; Methods); and include Anaphase Promoting Complex/Cyclosome (*Apc*) along with several other genes involved in centrosome organization (e.g. *Cenpe*, *Cep120*, *Camsap1*, *Numa1*). The first mitotic division in mammals is both significantly longer and more error prone compared to somatic mitotic events^27^. Furthermore, fertilization demarcates the beginning of a transition from multipolar acentrosomal division to the typical bipolar spindles organized by the centrosomes^28^. Critically, *Apc* activation is required for this reorganization and the dynamics of its activation underlies the prolonged first mitosis of mouse embryos^29^. Our results reveal differential translation efficiency as a key regulatory mechanism of this class of transcripts in the absence of any transcription.

When we compared 2-cell embryos to the zygote, we found a significant enrichment for RNA binding proteins among genes that had increased translational efficiency (Fisher’s exact test odds ratio 2.93; p-value 3.1 x 10^-10^; Supplementary Table 3). These include three genes that function as “readers’’ of N^6^-methyladenosine RNA modifications (*Hnrnpa2b1*, *Ythdf2*, and *Ythdc1;* Fig. 5B-C). Recent work has revealed all three of these genes to be required for successful early embryonic development^30–33^. Maternal depletion of *Ythdf2* in mice causes cytokinesis defects and arrest at the 2-cell stage ^31^. Similarly, reduced *Hnrnpa2b1* expression delayed embryonic development after the 4-cell stage^30^. Surprisingly, a detailed analysis of *Hnrnpa2b1* expression during preimplantation development had revealed negligible differences in RNA expression between the zygote and 2-cell mouse embryos, despite a dramatic increase in its protein abundance in 2-cell embryos^30^. Our analyses suggest that enhanced translation of *Hnrnpa2b1* is likely responsible for this observation. Intriguingly, while N^6^-methyladenosine “readers” had increased translation efficiency in 2-cell embryos, the key demethylase that removes N^6^-methyladenosine, Alkhb5^34^, was one of the most significantly downregulated genes in terms of translation efficiency (Fig. 5C; adjusted p-value 8.4 x 10^-08^). Taken together, our results reveal that translational regulation but not RNA expression as a shared mode of co-regulation of genes involved in N^6^-methyladenosine modification of RNAs.

### Ribosome occupancy is strongly associated with protein abundance during preimplantation mouse development

The proteome of the zygote is composed of maternally deposited proteins and those newly synthesized after fertilization^4^. However, the relative contribution of maternally deposited versus newly synthesized proteins to the zygotic proteome have not been possible to study at a transcriptome-wide scale in a mammalian system. Here, we applied Ribo-ITP to assess the contribution of translation in determining protein abundance^3^. In particular, protein abundance of 3,287 out of >5,000 transcripts detected in our single embryo ribosome profiling and RNA-Seq experiments had previously been quantified using mass spectrometry of ∼8000 embryos from each stage of mouse preimplantation development^3^.

We found that the zygotic proteome is only modestly correlated with RNA expression of the zygote (Spearman Rank Correlation 0.34; p-value < 2.2 x 10^-16^); in agreement with previous work that reported weak correlation between RNA expression and protein abundance^3, 35^. In contrast, zygotic protein abundance is significantly better correlated with zygotic ribosome occupancy than RNA expression (Spearman Rank Correlation 0.45 vs 0.34; p-value < 2.2 x 10^-16^; Figure 5D).

Critically, we discovered that ribosome occupancy of the GV-stage oocytes had the strongest relationship with the zygotic protein abundance (Fig. 5D; Spearman Rank Correlation 0.54, p-value < 2.2 x 10^-16^). Importantly, this key contribution is undetectable at the level of RNA expression as RNA abundance in GV-stage oocytes is much more weakly associated with the zygotic protein abundance (Spearman Rank Correlation 0.31; p-value < 2.2 x 10^-16^). Taken together, our results reveal that maternal translation is the predominant contributor to the zygotic proteome. This is a finding that eluded RNA-seq analyses and resolves the apparent paradox of low correlation between RNA and protein abundance at this critical stage in development.

The coupling of rapid degradation of the maternally deposited RNAs and onset of zygotic transcription fundamentally remodels the RNA contents of the developing embryo. Consequently, the 4-cell stage embryos have a very different RNA composition compared to 1- cell and 2-cell stage embryos. Intriguingly, neither ribosome occupancy nor RNA expression is positively correlated with protein abundance at the 4-cell or the 8-cell stage (Fig. 5E). Instead, we found that ribosome occupancy at the 4-cell and 8-cell stage embryos is much more strongly associated with the protein abundance at the morula stage. The association with protein abundance was significantly stronger for ribosome occupancy compared to RNA expression (Fig. 5E; Spearman Rank Correlation 0.66 vs 0.50 at 4-cell stage and 0.69 vs 0.53 at 8-cell stage, p-value < 2.2 x 10^-16^). These results suggest an unanticipated role of ribosome occupancy of the 4-cell and 8-cell stage embryos in determining morula protein abundance.

## Discussion

The development of ribosome profiling more than ten years ago was a landmark advance in our understanding of gene expression. However, unlike other sequencing-based assays such as RNA-Seq and ATAC-Seq that were swiftly adapted to analyze single cells within a few years of their initial development ^36, 37^, ribosome profiling remained limited to samples with a large number of cells, typically more than one million. In the conventional approach, RPFs are isolated through multiple days of hands-on experiments, involving ultracentrifugation, RNA extraction, and RNA size selection by polyacrylamide gel electrophoresis^15^. Together, these steps contribute to significant material loss, restricting the application of this technique to samples with abundant cellular inputs^38^. Consequently, many biological questions of importance remained beyond the scope of the conventional ribosome profiling in the last decade.

Here, we overcome this critical limitation by developing a novel microfluidic isotachophoresis approach, named RIBOsome profiling via IsoTachoPhoresis (Ribo-ITP). Unlike conventional techniques, Ribo-ITP maintains high yield recovery of RPFs and stringent RNA size selection even with picogram-level samples while simultaneously reducing sample processing time from several days to ∼30 min. The numerous innovations described in this study enable a significantly improved yield of recovery and previously unachieved stringency of RNA size selection in relation to previous on-chip ITP approaches^11, 12, 39, 40^. Compared with a recent study that applied single-cell RNA sequencing approaches in combination with an RNase digestion step^41^, Ribo-ITP specifically isolates RPFs from individual cells via a novel technology and consequently provide higher accuracy and coverage of measurements. Finally, the innovative use of 3’ ddC blocking modification and the optimized buffer exchange protocol in Ribo-ITP broaden the applicability of ITP-based sample purification to a variety of next generation sequencing-based analyses^42, 43^.

As an initial-proof-of-concept, we applied Ribo-ITP to study early stages of mouse embryonic development, where translational control of gene expression plays an imperative role^4, 44^. Our high coverage data enabled the first analysis of allele-specific ribosome occupancy and characterization of differential translation efficiency in preimplantation development. In particular, we discovered that APC/C and several components of the centrosome are translationally upregulated upon fertilization. The zygote needs to rely on maternally deposited mRNAs to initiate a mitotic program by remodeling the cellular environment transitioning away from meiotic divisions that proceed without centrosomes^45^. Hence, the initial preimplantation mitosis occurs under fundamentally different cellular conditions compared to somatic divisions^28^. Our results revealed translational upregulation as a previously unknown regulatory mechanism for key components involved in this transition including APC/C^29^.

Finally, we assessed the contribution of translation in determining the proteome of mouse preimplantation embryos. We discovered temporal dynamics that eluded previous RNA expression based analyses. Specifically, ribosome occupancy of GV-stage oocytes and not the zygote is the strongest predictor of zygotic protein abundance. This finding clarifies the previously observed poor correlation between RNA expression and protein abundance and suggests that proteins synthesized during oocyte maturation to be the predominant determinant of zygotic proteome. This study demonstrates the kind of new biological insights we can expect from the application of Ribo-ITP, which will help answer fundamental questions in translational control relevant to samples with limited input amounts including human biopsies, embryonic tissues, cancer stem cells and transient populations.

## Methods

### Polydimethylsiloxane (PDMS) chip fabrication

Molds were 3D-printed by Proto Labs with WaterShed XC 11122 at high resolution (Fig. S1). Reusable molds were assembled by taping 3D-printed molds to glass slides (5” by 4”; Ted Pella). Sylgard 184 PDMS monomer and curing agent (Ellsworth Adhesives 4019862) were mixed at a 10:1 (w/w) ratio. The mixture was degassed using a dessicator connected to a vacuum pump, poured over the mold, and degassed again until there were no air bubbles. The mold was incubated for at least 16 h at 50°C. Individual PDMS chips were cut along the lines that form the outer rectangle on the design in Fig. S1A. The 5 mm-diameter elution well, TE-, and LE-reservoirs were made with a biopsy punch (Fig. S1B). Prior to the plasma treatment, glass slides (4’’ by 3’’; Ted Pella) and the feature-side of the PDMS slabs were thoroughly cleaned with tape to remove any dust particles. PDMS chips and glass slides were plasma cleaned with a 115V Expanded Plasma Cleaner (Harrick Plasma) connected to a Dry Scroll Pump (Agilent) for 2 minutes at high RF level. The plasma-treated surfaces of the glass and PDMS slabs were immediately brought together to form a covalent bond. Bonded chips were heated at 80°C on a heat block for at least two hours to enhance bonding.

### PDMS chip preparation for Ribo-ITP experiments

To ensure clean, RNAse-free chips, we pre-treated the channels and reservoirs of the Ribo-ITP chip by sequential treatment with the following solutions: RNaseZap (100% concentrate), nuclease-free water, 1 M NaOH, nuclease-free water, 1 M HCl, nuclease-free water, 10% (w/v) benzophenone in acetone (for 10 minutes, replenishing channels as needed to avoid bubble accumulation), methanol, and 0.1% Triton X-100. The channel was completely dried after final treatment by fully vacuuming out any remaining liquid in the channel. After securing the chips to a ProteinSimple 302/365nm UV Transilluminator with tape, we added 10% polyacrylamide (PA) prepolymer mix (Supplementary Table 4) to the size-selection channel through the elution well. Similarly, 5% PA prepolymer mix was loaded into the extraction channel through branch channel 2. To catalyze the polymerization of polyacrylamide on chip, we used a photoactivatable azo-initiator, 2,2’-Azobis[2-methyl-N-(2-hydroxyethyl)propionamide] (Wako Chemicals VA-086), at 0.5% final concentration in the prepolymer mixes. UV-driven polymerization (365 nm wavelength) was performed for one minute followed by a 30 second break. This on/off UV cycle was repeated two more times for a total UV exposure time of 3 minutes. UV intensity was measured as ∼8.9 mW/cm^2^ using a G&R Labs Model 200 UV Light meter with a 365 nm probe. To avoid dehydration of the polyacrylamide gels after polymerization, we filled any open channels and reservoirs with storage buffer (Supplementary Table 4) until use. The chips were protected from light and used within six hours of preparation.

### Isotachophoresis setup

The prepared ITP chip was placed on a Dark Reader blue light transilluminator (Clare Chemical) and secured with tape. Storage buffer was removed from the channels and reservoirs using a vacuum. Leading electrolyte pluronic solution (LEp) and MOPS trailing electrolyte pluronic solution (TEp) (Supplementary Table 4) were kept on ice throughout the loading procedure. 200 µL pipet tips were kept at −20°C until the time of the experiment to facilitate manipulation of the pluronic-containing LEp and TEp solutions, which solidify within a minute above 4°C. 80 µL of LEp was loaded in LE reservoir 3, filling the reservoir to the top as well as the small section of the channel between the elution well and LE reservoir 3 (Fig. S1B). LE reservoir 2 was filled with 30 µL LEp, ensuring contact with the polyacrylamide gel present in branch channel 2. The elution well was filled with 20 µL of RB. Fluorescent marker oligonucleotides containing a 5’ ATTO fluorophore and 3’ ddC blocking modification (Supplementary Table 5) were added to the sample followed by dilution with sample dilution buffer (SDB). The mixture was loaded into the lysate channel through LE reservoir 1. Finally, LE reservoir 1 was filled with 30 µL LEp and 70 µl TEp was added to the TE reservoir. The negative electrode was placed in the TE reservoir and the positive electrode in the LE reservoir. Positive and negative electrodes were placed in LE reservoir 3 and the TE reservoir, respectively. A constant current of 300 mA with a maximum voltage of 1.1 kV (Keithley 2410 Sourcemeter) was applied to the channel. Once the trailing end of the fluorescent markers entered the 5% PA gel, the branch channel electrode–with a lower current output due to a 510 kΩ (Xikon) resistor on a custom circuit board–was manually applied in LE reservoir 1 for ∼10 seconds. When the leading edge of the longer fluorescent marker reached the end of the size-selection channel, the current was suspended. The elution reservoir was thoroughly washed twice with 30 µL nuclease-free water and refilled with 10 µL dephosphorylation buffer (Supplementary Table 4). Current was applied again until the shorter fluorescent marker began to enter the elution well. Finally, the purified sample with a 10 µL volume was collected from the elution well into a low-bind PCR tube and immediately stored at −80°C.

### Polyacrylamide gel electrophoresis and conventional extraction of RNA

Control inputs were prepared as a master mix then aliquoted. For gel extraction samples, input RNA was first processed using Qiagen miRNeasy Micro Kit per manufacturer’s instructions. RNAs were separated by electrophoresis using 15% TBE-Urea polyacrylamide gel (Invitrogen EC6885BOX). Gel slices were excised and crushed using sterile pestles, followed by soaking in gel extraction buffer (Supplementary Table 4) on dry ice for 30 minutes. Samples were then incubated overnight at room temperature, gently transferred on a tabletop shaker and protected from light. Residual gel pieces were removed by centrifugation for 1 minute at 21,130 x g through a Corning 0.22 μm sterile filter tube. The recovered eluate was precipitated overnight at −20 °C (300 mM sodium acetate pH 5.2, 5 mM MgCl_2_, 1.5 µL Glycoblue, 75% ethanol). Samples were pelleted by centrifugation at 4 °C for 1 hr at 21,130 x g.

### Gel imaging and quantification

To quantify yield, samples were run on a 15% TBE-Urea polyacrylamide gel and visualized using the fluorescent marker oligonucleotides or by SYBR Gold-staining. Specifically, gels were imaged using Typhoon FLA 9500 (GE Healthcare) with a 473 nm excitation wavelength and LPB filter compatible with ATTO488 fluorophore and SYBR gold-stain. For high resolution imaging, pixel size was minimized (10-25 µm) and PMT settings were optimized by using Typhoon’s scanning feature to avoid image oversaturation; typically resulting in a PMT value between 250 - 500 V. The images were analyzed using ImageJ software (NIH). Raw integrated density (RID) for background signal (RID_background_) was measured by quantifying average RIDs from representative blank areas. RID_background_ was normalized to account for the ratio of the target (A_sample_) to the background area (A_background_) such that

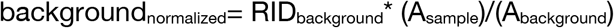

The normalized background value was subtracted from all samples to quantify normalized sample RID values. The percent yield was defined as the ratio of the normalized RID values to the mean of background-normalized input samples. For display purposes only, the contrast and brightness of some images were adjusted in ImageJ and exported as tiff files for figures.

### Yield comparison between on-chip method and conventional RNA extraction

Input controls and experimental samples were prepared with a final total amount of 40 ng, 20 ng, 2 ng, 400 pg, or 40 pg of ZR small RNA ladder (Zymo Research R1090) including 17, 21, 25, and 29 nt RNA oligonucleotides. Ribo-ITP was performed as described, with a final elution in 12 µL RB. Samples for gel extraction were first processed with the miRNeasy micro kit (Qiagen), followed by extraction using the crush & soak approach. Only the 25 nt and 29 nt bands were extracted. For 40 and 20 ng samples, fluorescent marker oligonucleotides were spiked into each sample and a final 15% TBE-Urea polyacrylamide gel was run as described above. Only the 25 nt and 29 nt bands were quantified to determine the final yield.

To quantify yield for the ultra-low input samples (2 ng, 400 pg, and 40 pg inputs), all experimental and input control samples were brought to 16 µL with nuclease-free water. Subsequently, 2 µL T4 polynucleotide kinase (PNK) buffer, 1 µL T4 PNK (NEB) and 1 µL ATP [γ-32P]- 3000Ci/mmol [10 mCi/mL] (Perkin Elmer NEG002A500UC) were added and incubated for 30 minutes at 37 °C. After incubation, unincorporated nucleotides were removed with the RNA Clean and Concentrator-5 kit (Zymo Research R1013) according to manufacturer’s instructions. RNA was eluted with 14 µL nuclease-free water, mixed with 2x denaturing gel loading dye (Supplementary Table 4) and denatured for 90 seconds at 80 °C. The samples were electrophoresed, then the gel was incubated in nuclease-free water for 5 minutes followed by a 30 minutes incubation in a 30% methanol and 5% glycerol solution. Both incubations were done on a rocking platform at room temperature. After the incubations, the gel was placed between pre-wetted cellophane sheets (Bio-Rad 1651779) and dried for 2 h in a GelAir drying system (Bio-Rad). The dried gel in cellophane was exposed for at least 12 h to a BAS-IP MS phosphor screen (GE 28956475). The phosphor screen was imaged with a Typhoon FLA 9500 (GE Healthcare) using 500 V PMT at 50 µM resolution. The image was visualized and quantified using ImageJ software only the 25 nt band was quantified as described above. All samples were processed in quadruplicate, with the exception of the Ribo-ITP sample with an RNA ladder input of 20 ng (n=3).

### Cell culture

Human K562 cells were grown in RPMI 1640 medium (Gibco) supplemented with 10% fetal bovine serum (Gibco) and 1% Pen-Strep (Gibco) and incubated at 37 °C with 5% CO_2_ to a density of ∼2.5 x 10^5^ cells per mL. Cells were regularly tested for Mycoplasma contamination. The authentication of K562 cell line was validated using STR profiling from ATCC.

### Size selection of purified RNA

To demonstrate the size selection capacity of our on-chip approach, we prepared an MNase-digested RNA sample from K562 cells. Briefly, 3 µL MNase (NEB) was added to a clarified K562 lysate from ∼5 M cells and digested for 30 minutes at 37 °C, followed by RNA extraction with the miRNeasy Micro kit (Qiagen) per manufacturer’s instructions. Ribo-ITP inputs contained 100 ng of the digested, purified RNA. Ribo-ITP was performed as described, with modifications to the collection method. Once the fluorescent marker band reached the interface of the 5% and 10% polyacrylamide gels, the current was suspended and RB was replaced with 12 µL of fresh RB. Ribo-ITP continued until the first fluorescent marker reached the edge of the elution well. The 12 µL of RB in the elution well was collected as Fraction 1 (F1). The well was washed twice with RB then refilled with 12 µL RB. Current was applied again until the front edge of the trailing fluorescent marker began to enter the elution well, and the 12 µL RB elution was collected as Fraction 2 (F2). The elution well was refilled with 12 µL RB and Ribo-ITP was continued for 2 minutes. The final 12 µL elution was collected as Fraction 3 (F3). Control inputs were prepared with the same amounts of bulk RNA and fluorescent markers, then brought to 12 µL with RB. Gel electrophoresis, imaging, and quantification were performed as described.

### Ribosome profiling sample preparation and monosome isolation

Approximately 10M K562 cells were pelleted, washed twice with PBS, and immediately flash-frozen in liquid nitrogen. Cells were lysed in 400 µL of cold lysis buffer (Supplementary Table 4) for 10 minutes on ice and pipetted to homogenize. The lysates were clarified by centrifugation at 1,300 x g for 10 minutes at 4 °C. Clarified supernatants were digested with 5 µL MNase (NEB M0247S) and incubated for 30 minutes at 37 °C. Digestions were stopped with 20 mM ribonucleoside vanadyl complex (NEB: S1402S). The samples were then loaded onto 20%-50% sucrose gradients and ultracentrifuged in a SW41 Ti swinging-bucket rotor (Beckman 331362) at 38,000 rpm for 2.5 h at 4 °C. The samples were fractionated using a Biocomp gradient fractionator. RNA was extracted from the monosome fractions with the miRNeasy Micro kit (Qiagen). One third of the eluate was electrophoresed through a 15% TBE-Urea polyacrylamide gel. The ribosome footprints of ∼17-35 nt were gel extracted using the crush-and-soak method as described. Final sample resuspension after ethanol precipitation was in 18 µL of nuclease-free water. The purified RNA was dephosphorylated with 1 µL of T4 polynucleotide kinase (NEB) in 1x T4 PNK buffer for 1 h at 37 °C. Dephosphorylated ribosome footprints were then ethanol precipitated (300 mM Sodium acetate, 2.5 volumes of ethanol, and 1.5 µL of GlycoBlue) overnight at −20 °C. Precipitated RNA was eluted in 10 µL nuclease-free water. The RNA was normalized to 350 ng in 6 µL of nuclease-free water before library preparation.

For 100-cell ribosome profiling experiments, K562 cells were pelleted, washed twice with PBS, and diluted to 100 cells in 5 µL of cold lysis buffer containing cycloheximide. MNase stock (2,000 gel units/ µL, NEB) was diluted 1:50 and 1 µL of the dilution was added to the samples. Digestion was performed for 30 minutes at 37°C in a thermal cycler with a heated lid. 1 µL EGTA was added to a final concentration of 10 mM in order to inhibit further digestion. Samples were placed on ice until processing through Ribo-ITP. Three replicates each were prepared for conventional ribosome profiling and Ribo-ITP.

### Mouse oocyte isolation

All experiments using mice by the Mouse Genetic Engineering Facility were approved by the Institutional Animal Care and Use Committee at the University of Texas at Austin. Oocytes were collected from superovulated C57BL/6J female mice as previously described ^23^. One hour after human Chorionic Gonadotropin (hCG) injection, the ovaries were placed in a 3 cm dish containing FHM medium (Cytospring, F1114), and Germinal vesicle (GV)-stage oocytes were released by scraping the surface of the ovaries with #5 Dumont forceps (Roboz). Meiosis II (MII)-stage oocytes were isolated from the oviducts approximately 14 h after hCG injection. Cumulus cells were removed from the oocytes by treatment with 1 mg/mL Hyaluronidase (Sigma H3884) in FHM medium. Both GV- and MII-stage oocytes were rinsed through three drops of FHM medium and then through three drops of 20 mg/mL BSA (Sigma A3311) in PBS (Hyclone SH30028.02). The oocytes were placed individually in 0.2 mL PCR tubes using a finely pulled glass pipette under a stereomicroscope and flash-frozen in liquid nitrogen. The liquid volume transferred with the oocytes was less than 0.5 µL.

### *In vitro* fertilization (IVF) using CAST/EiJ sperm

Sperm was frozen from CAST/EiJ male mice as described ^49^ and stored in liquid nitrogen. For *in vitro* fertilization, oocytes were isolated from C57BL/6J female mice approximately 15 h after hCG injection, and IVF was performed using thawed CAST/EiJ sperm^50^. 1-cell, 2-cell, 4-cell, and 8-cell embryos were collected 21.5, 39, 62, and 69 h after hCG injection, respectively. Fertilized oocytes were cultured overnight to the 2-cell stage in a 150 µL drop of HTF medium (Cytospring, mH0113). For development to the 4-cell and 8-cell stages, 2-cell embryos were cultured in KSOM medium (Cytospring, K0114). Embryos were placed individually into 0.2 mL PCR tubes and flash-frozen in liquid nitrogen. All samples were processed with Ribo-ITP within 48 h of collection.

A working lysis buffer solution was prepared by adding 1 µL of the MNase (NEB) [1:50 dilution] per 5 µL lysis buffer. To lyse the mouse samples, 6 µL of working lysis buffer was added directly to the frozen cell-containing droplet. Digestion was immediately performed for 30 minutes at 37 °C in a thermal cycler with a heated lid. 1 µL EGTA was added to a final concentration of 10 mM to inhibit further digestion. Samples were placed on ice until processing through Ribo-ITP.

### Ribosome profiling library preparation and sequencing

Conventional ribosome footprint libraries following monosome isolation (i.e. 350 ng RNA samples in 6 µL nuclease-free water) were generated using the Clontech SMARTer smRNA-Seq kit using 8 PCR cycles (Takara Bio). 30 µL of the PCR reaction was purified with AMPure XP beads (Beckman Coulter A63880) according to the manufacturer’s instructions and eluted with 30 µL of nuclease-free water. The final size selection was performed with the BluePippin system (Sage Science) using 3% dye-free agarose cassettes (Sage Science BDQ3010).

For 100-cell human K562 cells and all mouse samples, Ribo-ITP outputs were immediately processed through the D-Plex Small RNA-seq Kit (Diagenode C05030001) with minor modifications as detailed here. The dephosphorylation reaction was supplemented with 0.5 µL T4 PNK (NEB) and the reaction was incubated for 25 minutes. For reverse transcription, the template switching oligo (TSO) was diluted 1:2 in nuclease-free water. All 100-cell human samples and three of the MII-stage oocytes were processed using the single index (SI) module; while the other mouse samples were processed using the unique dual index (UDI) module. Half of the cDNA was amplified for 17 PCR cycles and a 1:4 dilution of the resulting library was assessed by Agilent Bioanalyzer High Sensitivity DNA kit. The concentrations of the target peaks were used to pool samples with approximately equimolar representation. AMPure XP bead cleanup (1.8x) was performed followed by size-selection using 3% agarose, dye-free gel cassettes with internal standards (Sage Science BDQ3010) on the BluePippin platform. Tight parameter settings of 173-207 bp range were used for samples prepared with the SDI module. Tight parameter settings of 183-217 bp range were used for samples prepared with the UDI module. Samples were sequenced on a Illumina NovaSeq 6000. For mouse samples, 5, 5, 5, 3, 3, and 4 biological replicates were used for GV, MII, 1-cell, 2-cell, 4-cell and 8-cell stages, respectively.

### Single cell and single embryo RNA sequencing (RNA-seq)

Total RNA sequencing libraries were prepared with Smart-seq3 V.3^51^, with modifications. Unfertilized mouse samples (GV, MII) and *in vitro* fertilized mouse samples (1, 2, 4, and 8-cell stage) were lysed and reverse transcribed as described. cDNA was pre-amplified with 13 PCR cycles and bead purified with AMPure XP (1.8x) with a final elution in 5 µL nuclease-free water. 1 µL of pre-amplified cDNA was assessed by Bioanalyzer High Sensitivity DNA kit to confirm successful pre-amplification and proper size profile. Another 1 µL was assessed on Qubit using the dsDNA HS assay to quantify the pre-amplified cDNA. Samples were diluted with nuclease-free water and normalized to 600 pg inputs [100 pg/µL] and subjected to tagmentation and post-tagmentation PCR. The tagmentation and subsequent PCR were scaled up 6x: precisely,, 600 pg pre-amplified cDNA was tagmented with 6 µL of Tagmentation Mix, 9 µL of Nextera Index primers were added, and 18 µL of Tagmentation PCR mix was used. 16 PCR cycles were performed followed by equivolume sample pooling (12 µL of each PCR product) and AMPureXP purification at a 1x ratio. The final library size distribution and concentration was assessed with the HS DNA Bioanalyzer. Sequencing was performed with Nova Seq 6000 with paired-end reads (using 100 cycle kits: 60+40). For GV, MII, 1-cell, 2-cell, 4-cell and 8-cell stages, 4, 4, 4, 4, 2 and 4 biological replicates were sequenced respectively.

### Computational processing of ribosome profiling data

Ribosome profiling data were processed using RiboFlow ^52^. We extracted the first 12 nucleotides from the 5’ end of the reads using UMI-tools^53^ version 1.1.1 with the following parameters: “umi_tools extract -p ″\(?P<UMI_1>.{12})(?P<DISCARD_1>.{4}).+$″ --extract-method=regex”. The four nucleotides downstream of the UMIs are discarded as they are incorporated during the reverse transcription step. Conventional ribosome profiling samples did not include UMIs.

Next we clipped the 3’ adapter AAAAAAAAAACAAAAAAAAAA, from the Ribo-ITP data, using cutadapt ^54^ version 1.18 with the parameters “-a AAAAAAAAAACAAAAAAAAAA -- overlap=4 --trimmed-only”. For conventional ribosome profiling data, we removed the poly-A tails and the first three nucleotides of the reads using “cutadapt -u 3 -a AAAAAAAAAA -- overlap=4 --trimmed-only”.

After UMI extraction and adapter trimming, reads were aligned to ribosomal and transfer RNAs using Bowtie2 ^55^ version 2.3.4.3. The unaligned reads were mapped to a manually-curated transcriptome. We retained alignments with mapping quality greater than 2 followed by deduplication using UMI-tools when applicable. In deduplication of external libraries without UMIs, a set of reads with the same length that are mapped to an identical nucleotide position are collapsed into a single read. As the last step, .ribo files are created using RiboPy^52^ version 0.0.1. All subsequent analyses used ribosome footprints that were 29-35 nucleotides in length.

For analyses involving nucleotide resolution data, we determined the A-site offset for each ribosome footprint length using translation stop site metagene plots. Specifically, for each read length, we identified the highest peak upstream of the translation stop site and used the distance to the annotated stop site as the offset.

### Computational processing of RNA sequencing data

5’ adapter sequence “ATTGCGCAATG” was clipped from the first read in the pair using cutadapt ^54^ version 1.18. Clipped reads shorter than 8 nucleotides were removed using: “cutadapt -j 4 --trimmed-only -m 8 -g ATTGCGCAATG”. We then extracted the next 8 nucleotides corresponding to the UMIs from the first read in the pair and appended them to the headers (of FASTQ files) of both read pairs using UMI-tools with the following parameters: “umi_tools extract --bc-pattern NNNNNNNN”. After UMI extraction, we used the second read in the pair (40 nt) for all subsequent analyses.

After filtering out reads aligning to a reference of ribosomal and tRNAs, the remaining reads were aligned to a transcriptome reference where SNPs were masked with Ns (see the next section for details); thereafter, we retained only the alignments with mapping quality greater than 2. We then collapsed reads that align to the same transcript using their respective UMIs: “umi_tools dedup --per-contig --per-gene”. For each transcript, we counted the number of reads aligning to the coding sequence. We used bowtie2 ^55^ for all alignments and samtools ^56^ version 1.11 for processing BAM files.

### Comparison with polysome profiling

The transcripts with validated changes in polysomal association between GV- and MII-stage oocytes were obtained from Supplemental Figures S2 and S3 of Chen et al.^23^. Of the 29 genes with qrt-PCR validated changes in polysomal association, 28 had the reported direction of effect when comparing the mean of the centered log-ratio (CLR)^57^ across the replicates. Specifically, Let *M* be the geometric mean of all the genes with non-zero counts and let *g* be the raw counts for a specific gene. Then, CLR of *g* is computed as 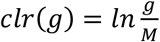.

### Allele-specific ribosome occupancy and RNA expression analysis

A list of strain-specific single nucleotide polymorphisms (SNPs) was obtained in VCF format from https://github.com/sandberg-lab/Smart-seq3/blob/master/allele_level_expression/CAST.SNPs.validated.vcf.gz ^51^. We extracted 210,004 distinct SNPs that overlapped with transcript annotations. To avoid alignment biases, we modified our transcriptome reference sequences by masking SNP positions with Ns. Mouse sequencing data were aligned to this masked transcriptome reference.

For allele-specific analyses reported in Fig. 4A, we considered the 85,339 SNPs within the coding sequences of transcripts. Given that transcripts in oocytes should solely contain maternal SNPs, we used the data from the MII-stage oocytes to construct a simple error correction model. Specifically, 2.67% and 0.40% of reads contained non-maternal sequences in ribosome profiling and RNA-Seq experiments, respectively. These values were used as estimates of the sequencing error percentage (*error*).

We define the paternal ratio as 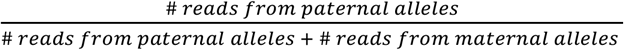. For 1-cell to 8-cell embryos, we then calculated the error-corrected paternal ratio, *paternal_corrected_* as:

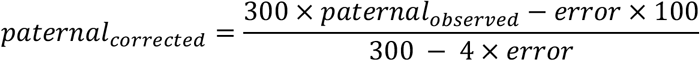

where, *paternal*_observed_ is the uncorrected percentage. We derived this equation from the below model under the assumption that sequencing errors are random:

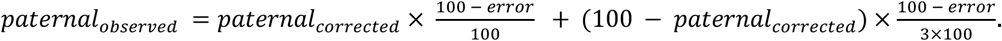

For each embryonic stage, to identify the transcripts whose paternal ratios are significantly different in ribosome profiling compared to RNA-Seq, we first aggregated all SNP-containing reads for each transcript across replicates. We retained transcripts with more than 10 reads in both ribosome profiling and RNA-seq experiments including at least three maternal and paternal reads. We used a two-sample test for the equality of proportions with continuity correction (prop.test in R; see Chapter 3 of ^58^ for details).

Transcripts with 95% confidence intervals of difference in paternal ratios (derived from the test for the equality of proportions), overlapping with the interval [-0.05, 0.05], are filtered out. After adjusting the p-values using the false discovery rate method, we retained the transcripts with adjusted p-values less than 0.2. We further removed transcripts with paternal reads in the MII-stage as these likely indicate positions that are prone to alignment errors. As the final step, we applied bootstrapping to establish robustness of the conclusions. Specifically, we randomly sampled replicates with replacement and repeated the statistical testing procedure described above. 24 transcripts with a false discovery rate less than 0.2 in at least 66 out of 100 bootstrap samples were deemed as having differential allelic ratios.

### Differential expression and translation efficiency analysis

Reads that align to coding regions were extracted for all experiments. To determine Transcripts with the highest variability in ribosome occupancy across developmental stages, a variance-stabilizing transformation (VST), as described in ^59^, was applied to centered log-ratio of ribosome occupancies (“FindVariableFeatures” function with the selection method “vst” in the Seurat package v4 ^60^). Using the threshold “vst.variance.standardized” > 4.8, we obtained 50 genes.

For every pair of consecutive developmental stages, differential RNA expression and translation efficiency was determined using DESeq2^48^. For differential translation efficiency calculation, we used the interaction term between the developmental stage and measurement modality (ribosome profiling or RNA-Seq). Default parameters were used for read count normalization and estimation of gene-specific dispersion. Effect size moderation was carried out using the approximate posterior estimation for a generalized linear model ^61^. The adjusted p-value cutoff was set to 0.01 to determine a set of transcripts with significant changes in RNA expression and translation efficiency. Gene set enrichment analyses for gene ontology terms were carried out using FuncAssociate (http://llama.mshri.on.ca/funcassociate/) with default settings^47^.

### Proteomics data and comparison with RNA-Seq and ribosome profiling

TMT-labeling based proteomics abundance data for 1-cell to morula stage embryos was obtained from Gao et al.^3^. 3287 proteins had measurements in all three modalities and were used in further analysis. Ribosome occupancy and RNA expression were converted to read density by dividing the read counts by the length of the coding region of each transcript. These values were normalized using a centered log ratio transformation as implemented in Seurat v4^60^. The similarity between RNA expression, ribosome occupancy and protein abundance was measured using rank correlation with Spearman’s correction^62, 63^. The measurement reliability for each modality was estimated using replicate to replicate correlation coefficients (0.71 for ribosome profiling, 0.79 for RNA-seq and 0.8 for mass spectrometry^3^).

### Weighted Transcript Region Length Distribution

For the transcript regions, 5’ UTR, CDS and 3’ UTR, the distribution of weighted region lengths was calculated as follows. First, for each transcript, we determined the ratios of region lengths to the transcript length. Next, we multiplied these ratios with the number of ribosome occupancies in the transcript, giving us weighted ratios of the regions. Then, for each region, we calculated the sum of their weighted ratios across transcripts. Finally, let *w*_5, *UTR*_, *W_CDS_* and *W*_3, *UTR*_ be the weighted sums of the regions 5’ UTR, CDS and 3’ UTR, respectively. For a region r, the weighted length percentage of r is 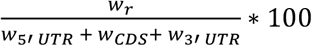.

## Acknowledgements

We would like to thank Dr. William Shawlot and the UT Austin Mouse Genetic Engineering Facility for their contributions. The authors would also like to thank Dr. Arlen Johnson, Dr. Steve Vokes, Dr. Basar Cenik, Dr. Shilpa Rao, Ian Hoskins and Dr. Elif Sarinay for their critical evaluation of the manuscript and contributions to the writing.

Certain commercial equipment, instruments or materials are identified in this paper in order to specify the experimental procedure adequately. Such identification is not intended to imply recommendation or endorsement by the National Institute of Standards and Technology (NIST), nor is it intended to imply that the materials or equipment identified are necessarily the best available for the purpose.

## Author contributions

T.T.: Methodology, validation, formal analysis, visualization, writing—original draft, writing—review and editing. H.O.: Data curation, investigation, validation, formal analysis, software, visualization, writing—original draft, writing—review and editing. C.H.: conceptualization, methodology, formal analysis, writing—original draft, writing—review and editing. A.S.: Visualization, methodology D.C.: Investigation, validation M.S: Conceptualization, project administration, writing— review and editing. C.C.: Conceptualization, data curation, formal analysis, software, supervision, funding acquisition, investigation, methodology, project administration, visualization, writing—original draft, writing—review and editing.

## Data and materials availability

Sequencing files for ribosome profiling and RNA-seq experiments, together with additional supplemental files, have been deposited to NCBI GEO database and accession number will be updated upon acceptance. The code used in the study is available at https://github.com/CenikLab/ribo-itp_paper.

---

**Extended Data Figure 1.**
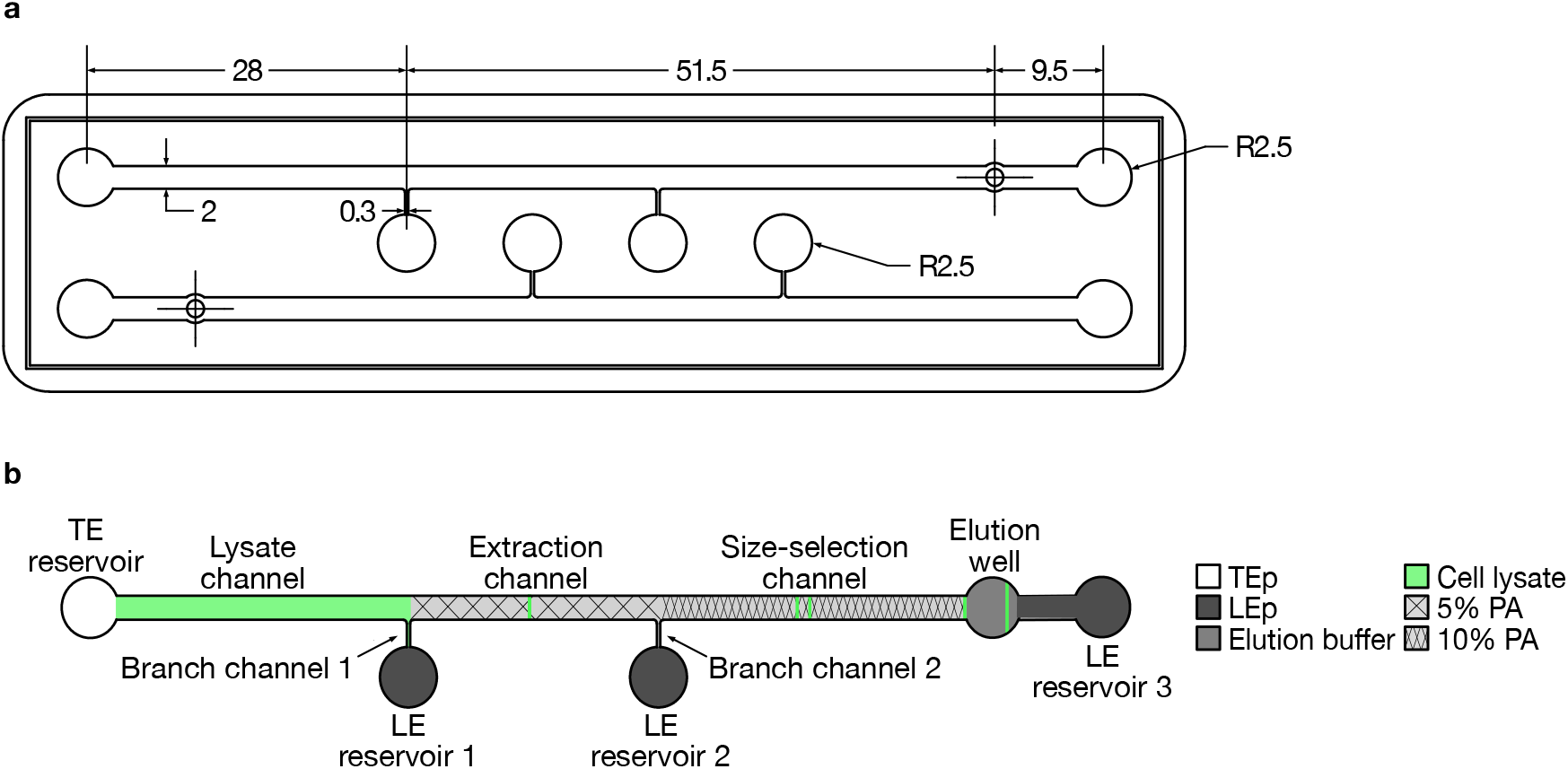
Channel design and dimensions. **a** The top view of the ITP chip layout designed with SOLIDWORKS (units in mm). The design was 3-D printed to be used as a mold for microchannels. The thickness of the channel features was 375 µm and that of the rectangular base was 1.5 mm. Linear tapering was applied from the rectangular base to the outer edge (rounded rectangle). **b** Microfluidic device setup; indicating lysate, extraction, and size-selection channels; trailing electrolyte (aTE) and leading electrolyte (LE) reservoirs (1-3); and elution well. Buffers corresponding to each channel and reservoir are color coded. Marker oligonucleotide fluorescence is denoted by green.

**Extended Data Figure 2.**
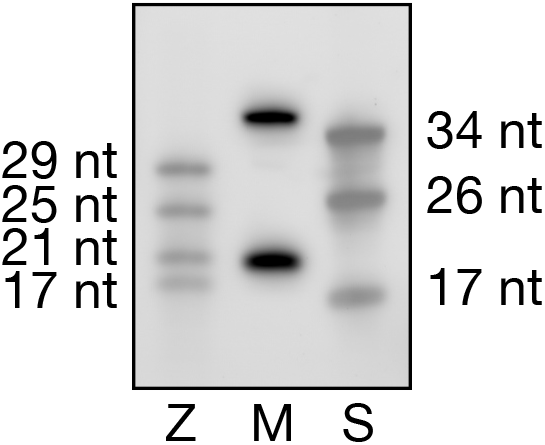
Verification of encapsulation of typical RPF size range by fluorescent DNA marker oligonucleotides. The gel image displays the relative mobilities of small RNAs and modified fluorescent markers used in Ribo-ITP. The Zymo Research R1090 small RNA ladder (Z) and synthetic RNA oligonucleotides (S) exemplify potential fragment lengths generated by MNase digestion. 19 nt and 36 nt fluorescent DNA oligonucleotide markers (M) are used in Ribo-ITP experiments.

**Extended Data Figure 3.**
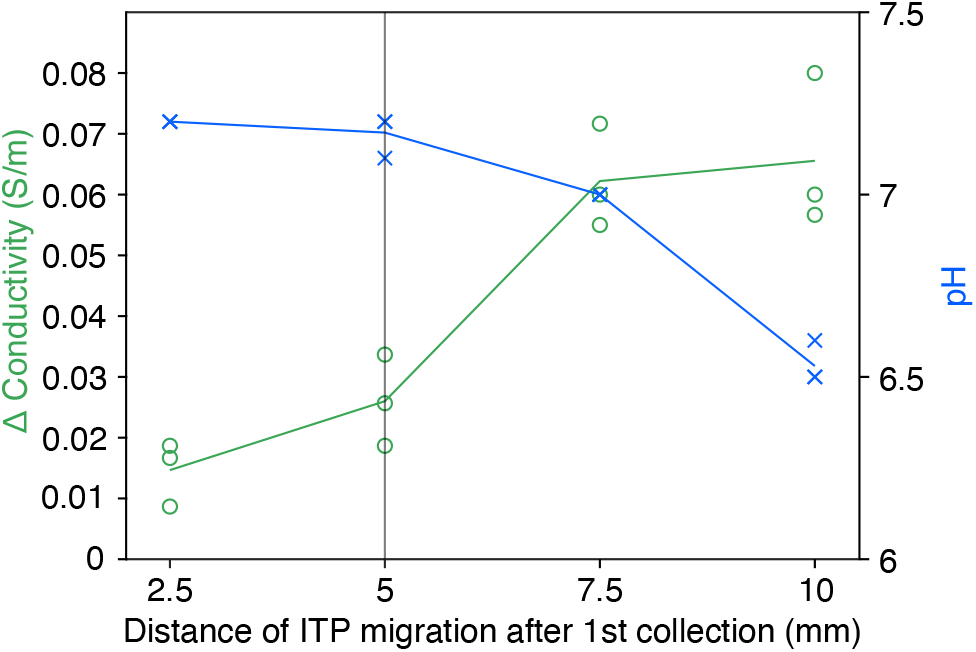
Conductivity and pH measurements of dephosphorylation buffer subjected to Ribo-ITP. The first step of library preparation in Ribo-ITP is 3’ dephosphorylation. Given that pH is the critical factor determining dephosphorylation efficiency45, we determined the impact of Ribo-ITP collection on the conductivity and pH of the dephosphorylation buffer. We found negligible pH change (right axis, blue) and only a 11.0% ± 1.83% (SEM) change in conductivity (left axis, green) for the collection distance (5mm, denoted by the vertical line) used in Ribo-ITP.

**Extended Data Figure 4.**
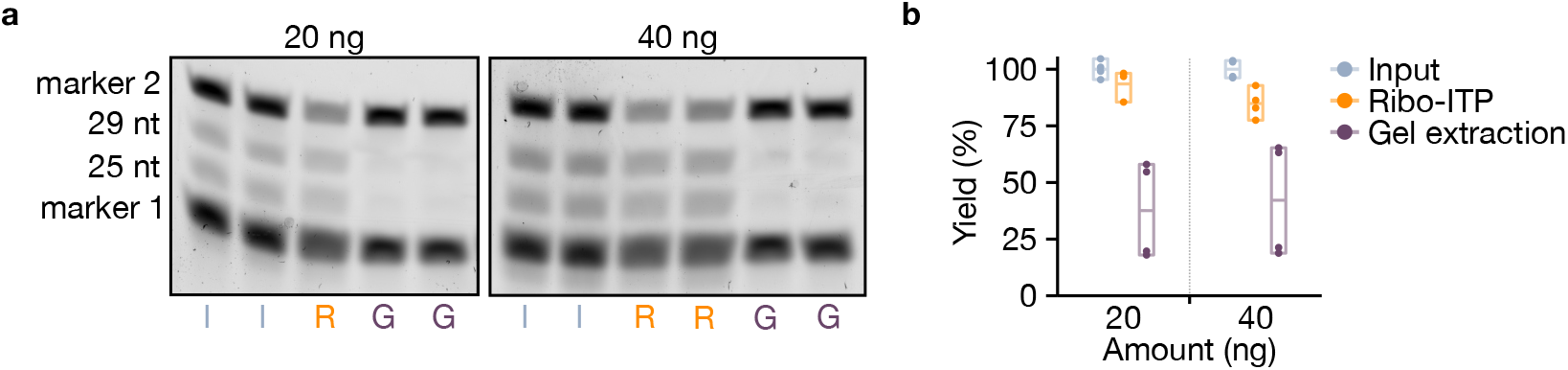
Yield comparison between Ribo-ITP and conventional gel extraction. **a,** Representative gel images of control inputs (I), Ribo-ITP elutions (R), and gel extraction (G) samples. Four RNA species (17, 21, 25, and 29 nt) were used with total inputs of 20 and 40 ng. Fluorescent marker oligonucleotides were spiked into control and gel extraction samples prior to gel visualization. **b,** Gel image quantification of control inputs (gray), Ribo-ITP elutions (orange), and gel extraction (purple) samples. Minimum, maximum, and average values are represented by the box and the horizontal bar. Only the 25 and 29 nt RNA marker bands were quantified for the yield calculation.

**Extended Data Figure 5.**
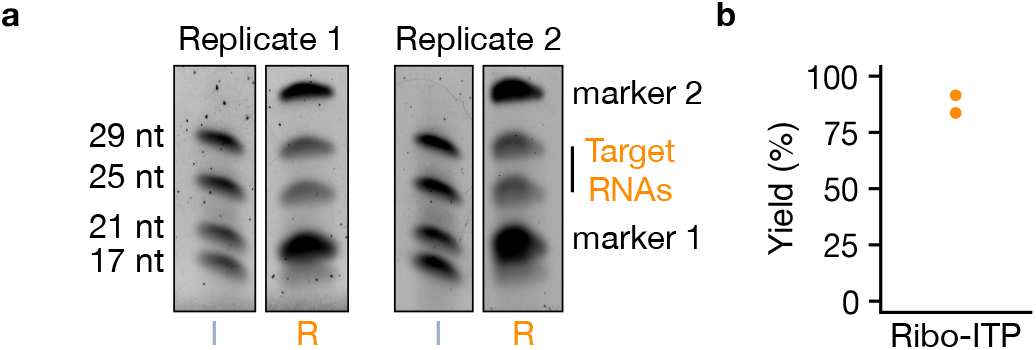
Ribo-ITP enables efficient RNA extraction from cell lysates. **a,** Inputs (I, gray) were prepared by adding 40 ng of RNA to lysates from ∼1,000 K562 cells. The RNA consisted of four species ranging from 17 to 29 nt in length. Fluorescent marker DNAs were added to Ribo-ITP samples (R) in addition to EGTA (10 mM). RNA extraction and isolation was done with Ribo-ITP followed by visualization using gel electrophoresis. **b,** Yield of the 25 and 29 nt RNAs was quantified and plotted for two replicates (84% and 91%, respectively).

**Extended Data Figure 6.**
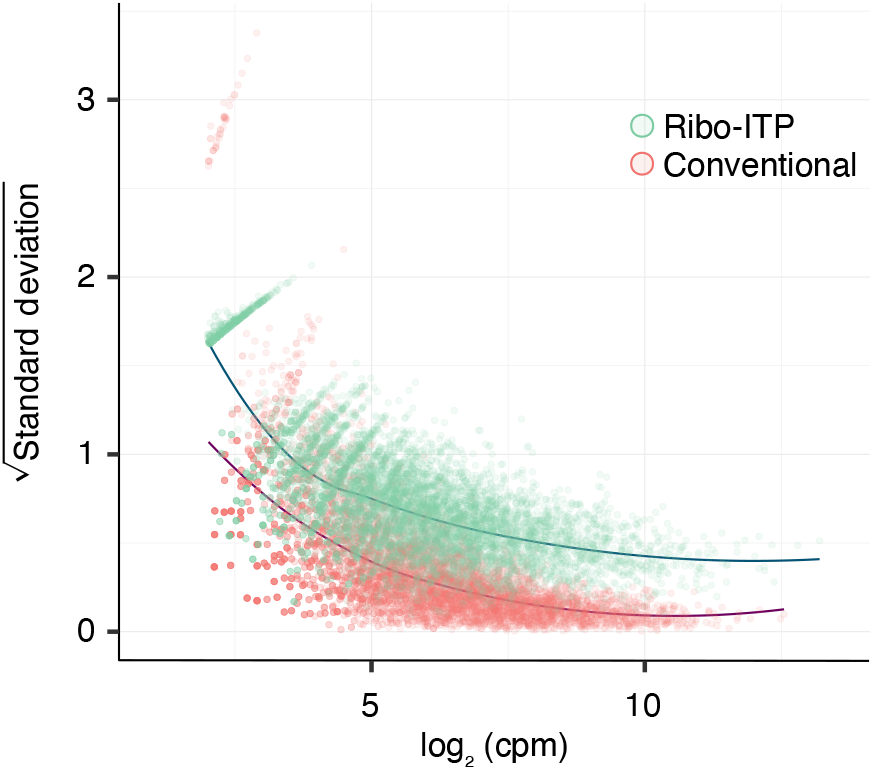
Mean to dispersion relationship for Ribo-ITP and conventional ribosome profiling. Transcripts with at least one count per million (cpm) in at least two out of three replicates were selected. Log2 of mean cpm (x-axis) was plotted against the square root of the standard deviation of cpm values (y-axis). Green (Ribo-ITP from ∼100 cells) and red (conventional ribosome profiling from ∼10M cells) points represent individual transcripts with mean log2(cpm) greater than two.

**Extended Data Figure 7.**
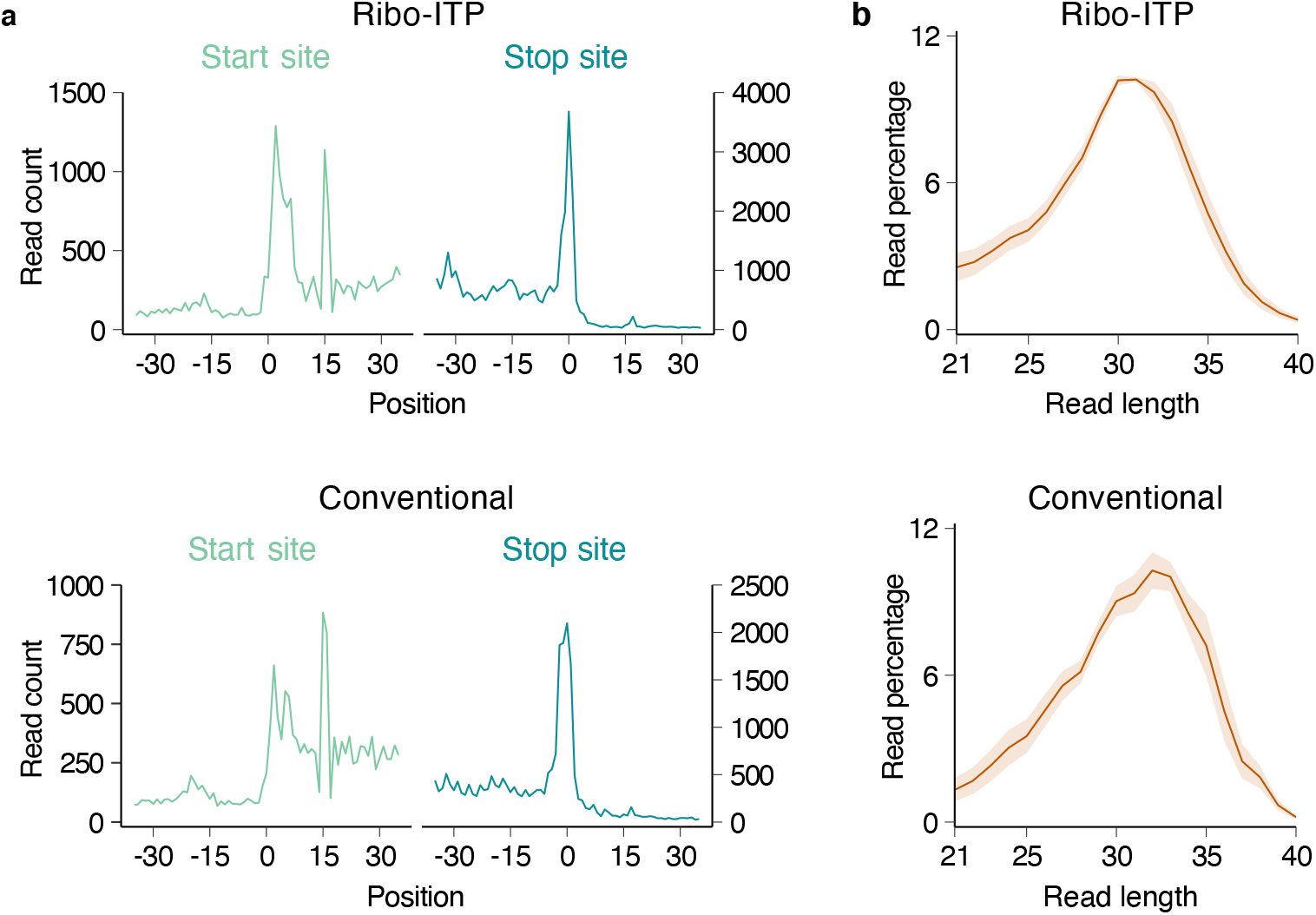
Metagene plots and footprint length distribution for Ribo-ITP from ∼100 cells compared to conventional ribosome profiling from ∼10M cells. **a,** Metagene plots of Ribo-ITP versus conventional ribosome profiling in human K562 cells were shown. Position 0 corresponds to the start (left, light green) or stop (right, dark green) site. Aligned positions of ribosome footprints were adjusted according to their A-site offsets. One representative replicate for each method is plotted. **b,** The mean percentage of specific read lengths among the total mapped reads is plotted. Ribbons around the lines represent standard error of the mean.

**Extended Data Figure 8.**
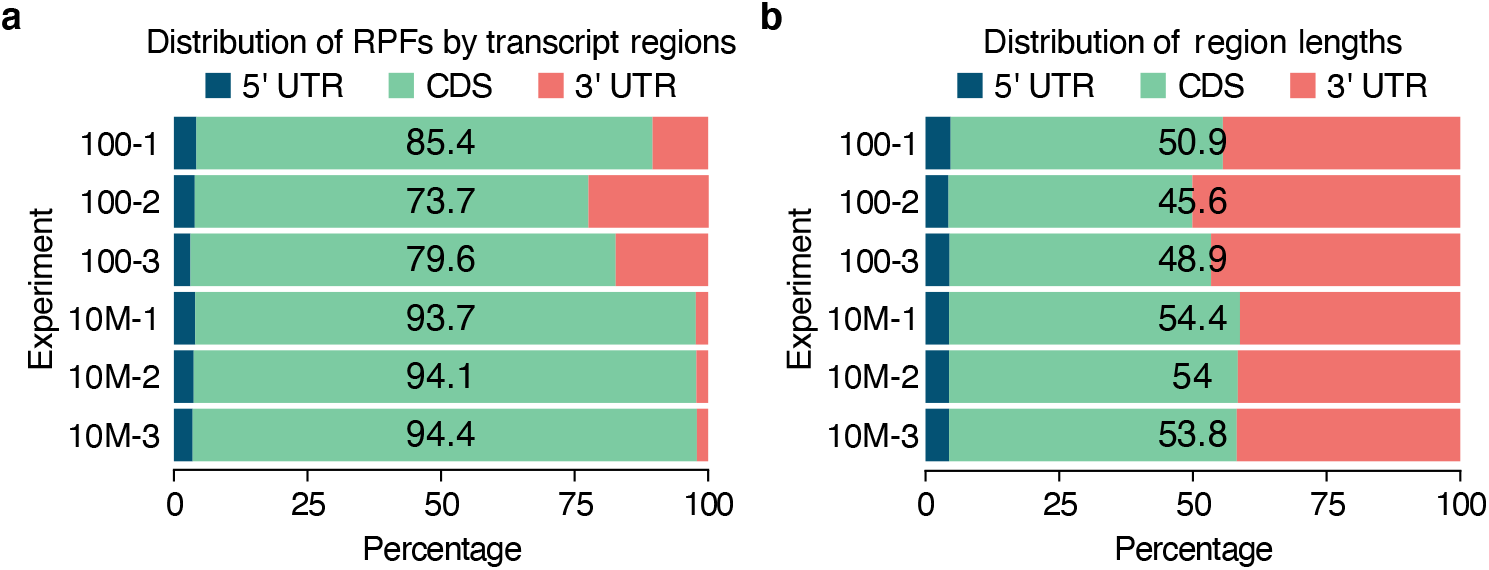
Region Counts for Ribo-ITP from ∼100 cells compared to conventional ribosome profiling from ∼10M cells. **a,** Percentages of ribosome profiling reads aligning to different transcript regions are plotted. The percentage of reads mapped to the CDS is indicated for each experiment. **b,** For each transcript, we multiplied its region length by the total number of ribosome footprints for the given experiment. The plotted overall percentage corresponds to the sum of these weighted counts across transcripts.

**Extended Data Figure 9.**
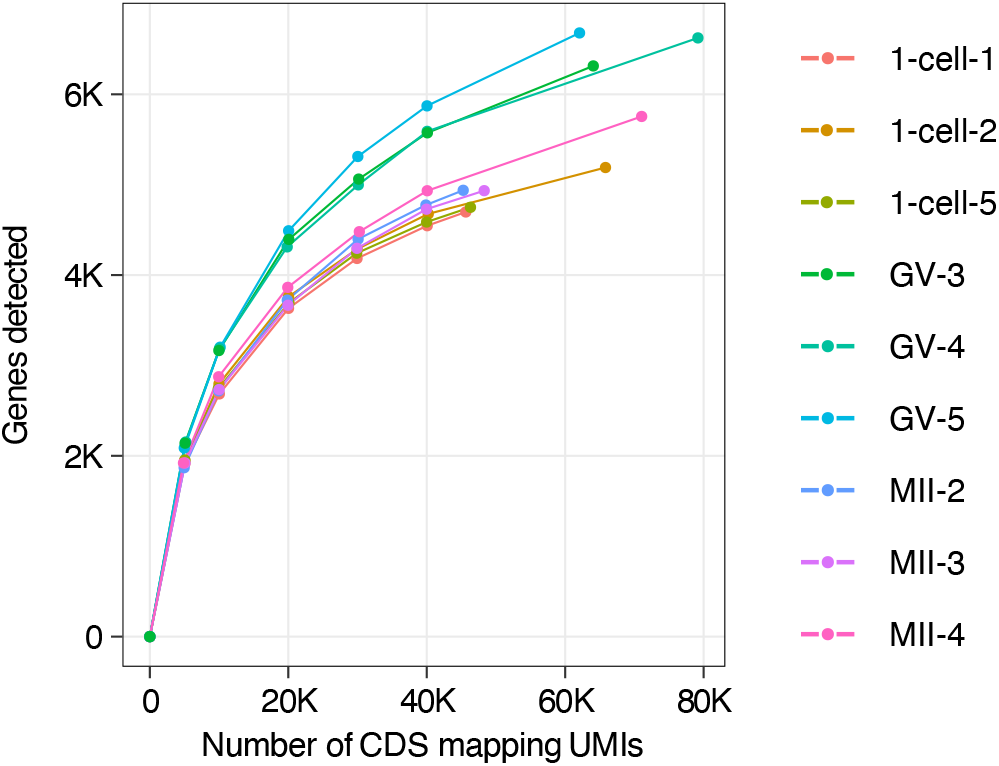
Number of genes detected as a function of CDS mapping UMIs. The number of reads (x-axis) is plotted against the number of detected genes (y-axis). Three representative replicates are shown for each stage of development used for single cell ribosome profiling experiments. The total number of detected genes using all CDS mapping reads is indicated along with genes detected with 5k, 10k, 20k, 30k and 40k sub-sampled coding regions mapping UMIs.

**Extended Data Figure 10.**
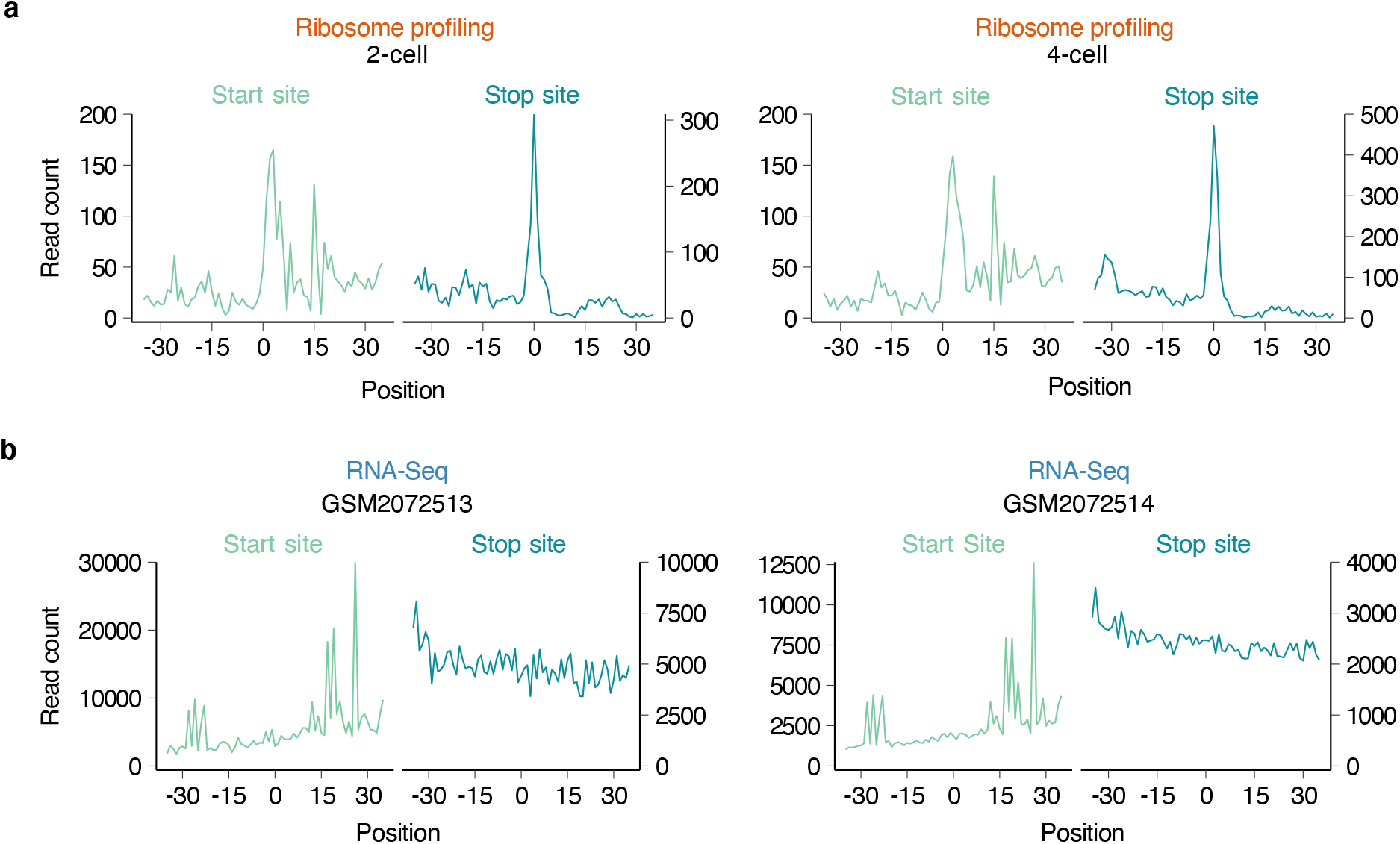
Metagene plots for mouse Ribo-ITP and RNA-Seq experiments. **a,** Metagene plots of translation start and stop sites from a representative Ribo-ITP experiment using a 2-cell or a 4-cell stage mouse embryo. Start sites (light green, left) and stop sites (dark green, right) are at position 0 on the x-axis. Positions of the aligned reads are adjusted according to their A-site offsets. **b,** Metagene plots of translation start and stop sites from RNA-Seq data. GEO accession numbers of the experiments are indicated on the plots. In contrast to the ribosome profiling data, there is no detectable peak is observed at translation start or stop sites.

**Extended Data Figure 11.**
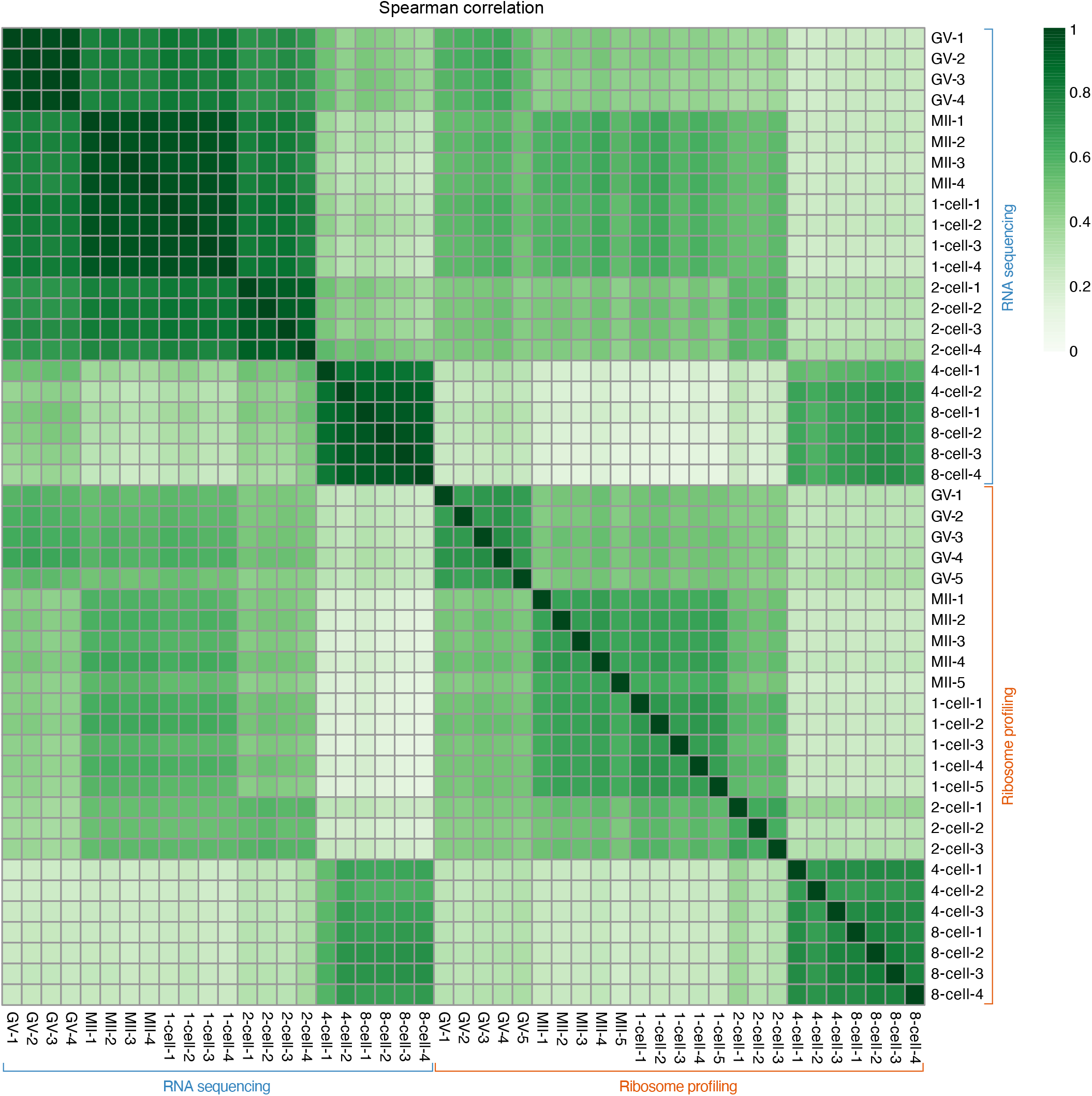
Pairwise correlation of read counts from Ribo-ITP and RNA-Seq experiments. CDS-mapping read counts from each transcript were used to compute the Spearman correlation coefficient. Ribo-ITP (orange) and RNA-Seq (blue) experiments are ordered by developmental stage. The colors indicate the strength of the correlation.

**Extended Data Figure 12.**
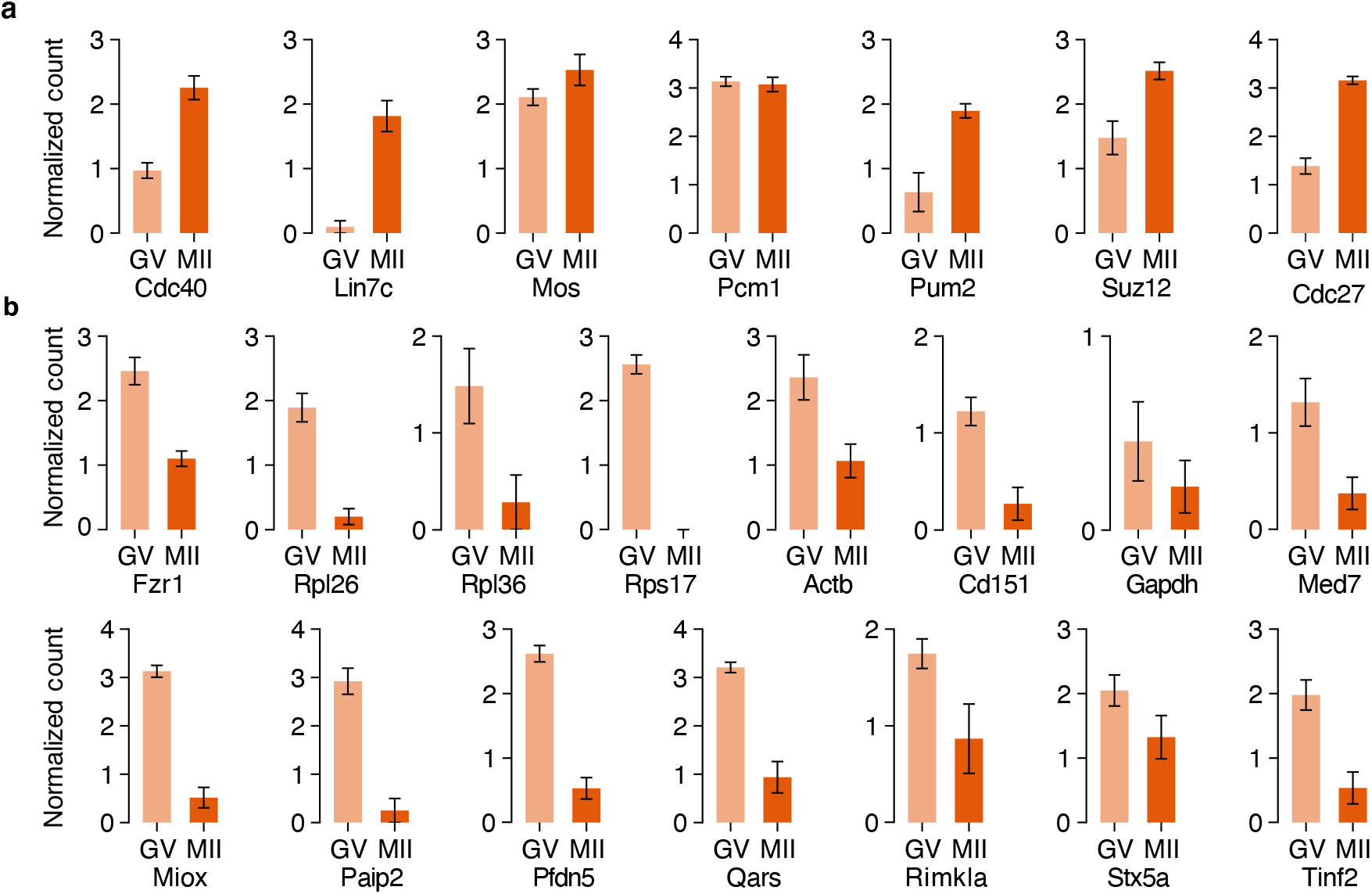
Ribosome occupancy in GV- and MII-stages of transcripts with previously identified differential polysome association. **a,** The mean of centered log-ratio of ribosome occupancy (y-axis) was plotted along with the standard error of the mean. These transcripts were identified as having increased polysome association in the MII-stage compared to GV-stage^24^. **b,** Similar to panel a with the exception that these transcripts were found to display decreased polysome association in the MII-stage compared to the GV-stage^24^.

**Extended Data Figure 13.**
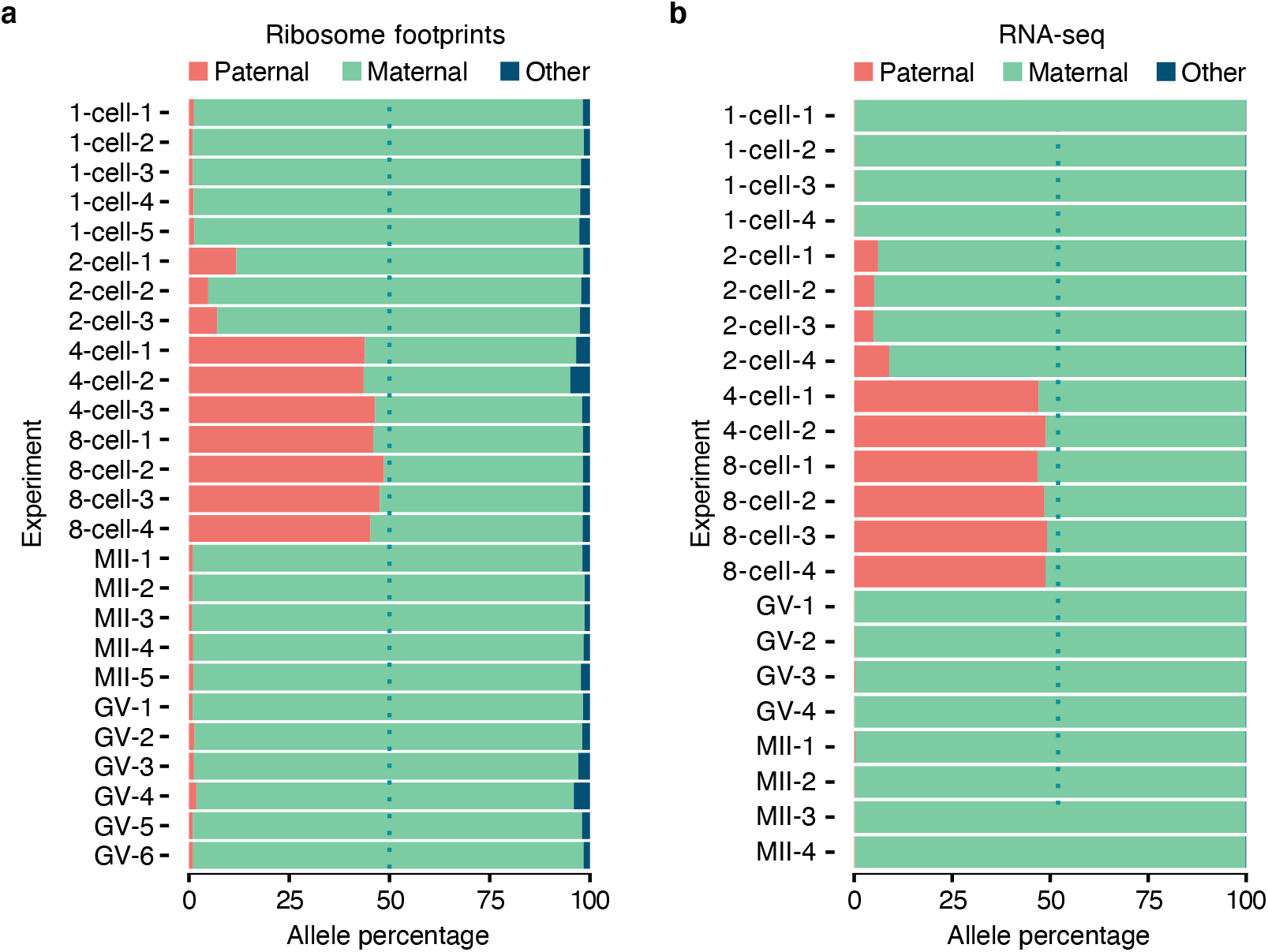
Distribution of sequencing reads that overlap strain-specific SNPs. **a,** Ribosome footprints that overlap strain-specific SNPs were used to determine the percentage of reads that match the maternal (green) and paternal (red) allele. A small percentage of reads differed from either allele and are labeled as “other” (dark blue). **b,** Reads from RNA-seq experiments were used as in panel a.

**Extended Data Figure 14.**
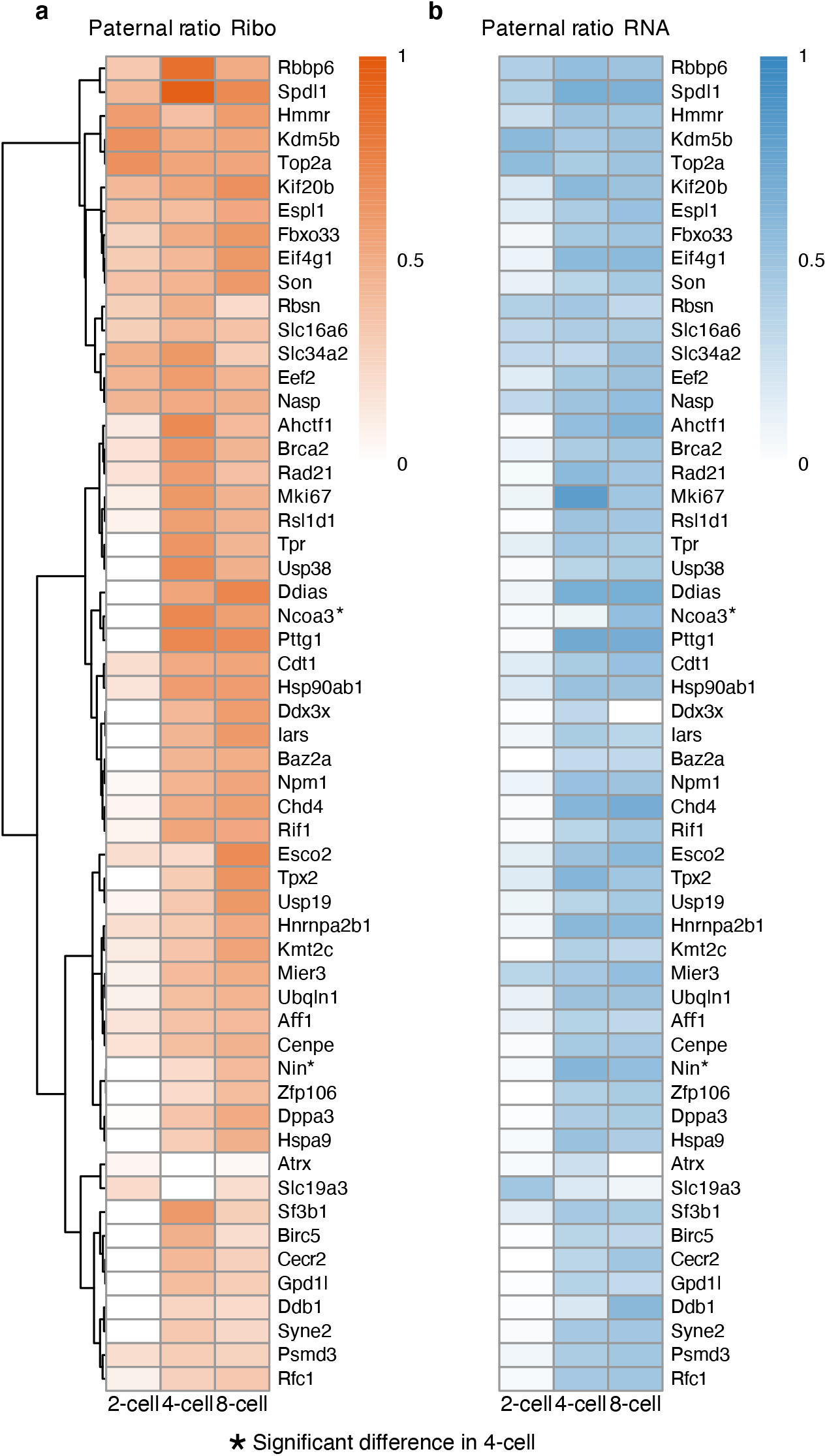
Allele-specific ribosome occupancy and RNA expression of genes with the highest number of informative reads. **a,** The percentage of ribosome footprints that originate from the paternal allele was visualized for genes with at least 10 allele-specific reads at each stage. **b,** The percentage of RNA sequencing reads that were assigned to the paternal allele was plotted for the set of genes in panel a.

**Extended Data Figure 15.**
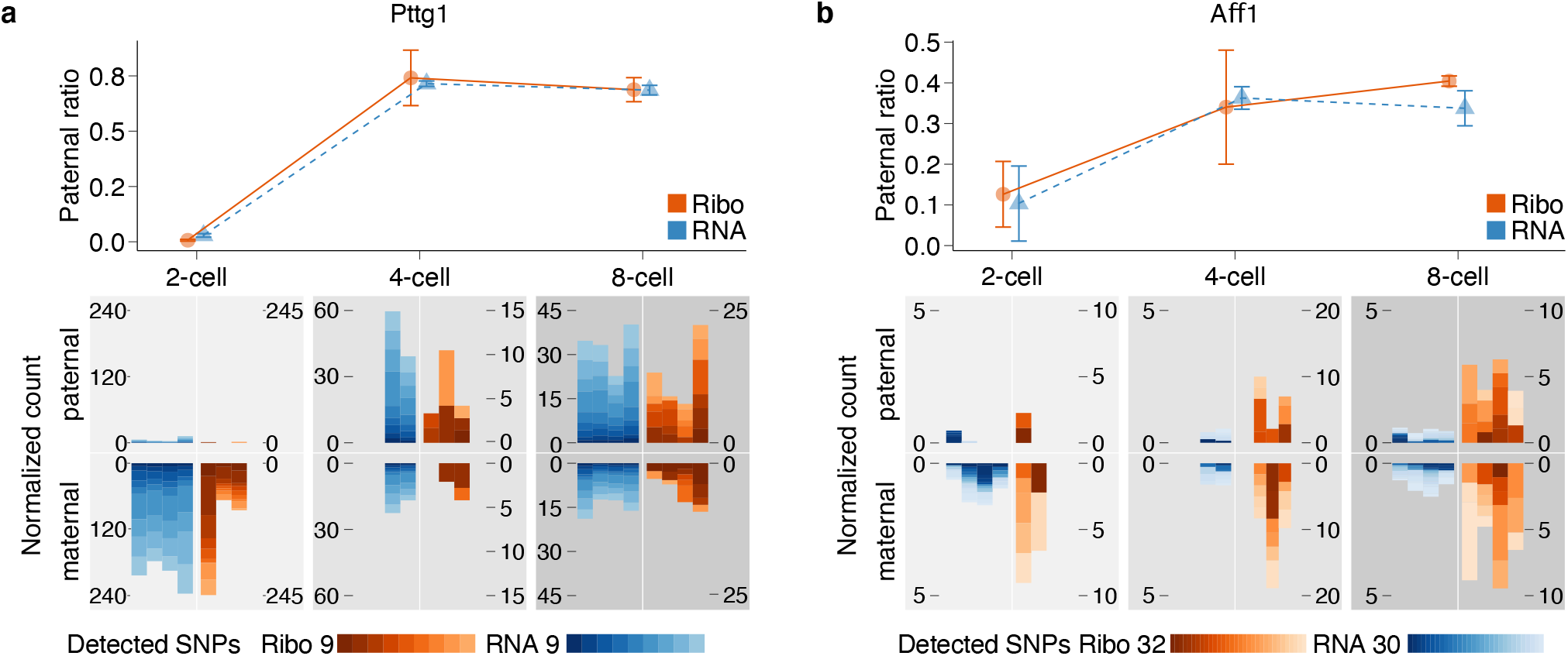
Representative genes with no allele-specific differences between RNA expression and ribosome occupancy. **a,b** Line-plots (top) indicate the percentage of paternal reads (y-axis) in RNA-Seq and Ribo-ITP experiments. The reads are combined across replicates and error bars indicate standard error of the mean of paternal ratios. At the bottom, maternal and paternal reads counts per 10k are plotted for all individual replicates and SNPs. The total number of detected coding SNPs and their corresponding colors are shown with color scales. Each vertical bar corresponds to a replicate experiment.

**Extended Data Figure 16.**
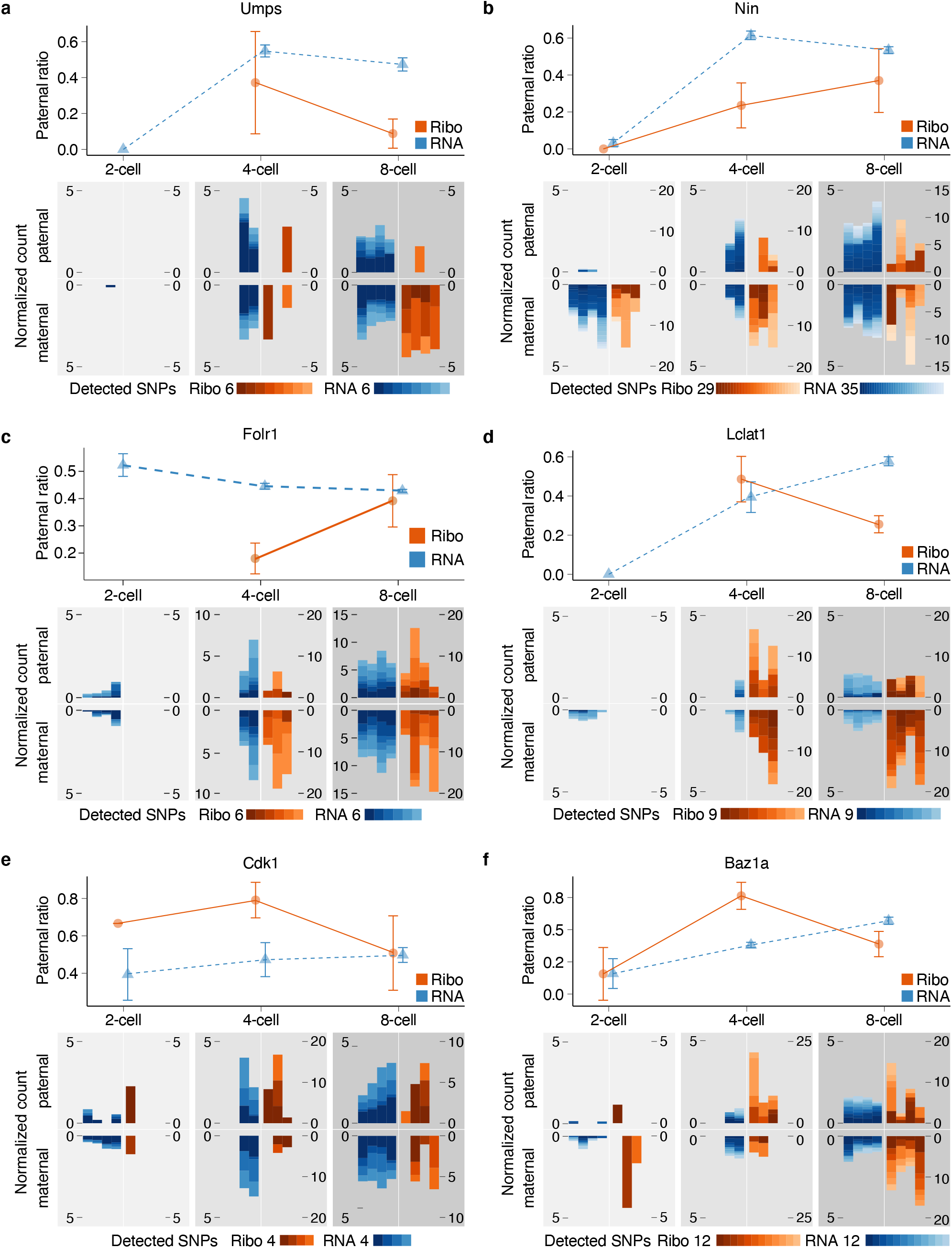
Representative genes with allele-specific difference in ribosome occupancy and RNA expression. Similar to S15. **a,** Representative gene from cluster II with allele-specific ribosome occupancy bias (e.g of maternal bias) **b, c,** Representative genes from cluster III with delayed engagement of ribosomes in allele- and stage-specific manner i.e. parental RNA expression is observed at or before the 4-cell stage, yet these RNAs predominantly engage with ribosomes only in the 8-cell stage **d,** Representative genes from cluster IV with differential ribosome occupancy compared to RNA expression in 8 -cell stage and **e, f,** in 4- cell stage.

**Extended Data Figure 17.**
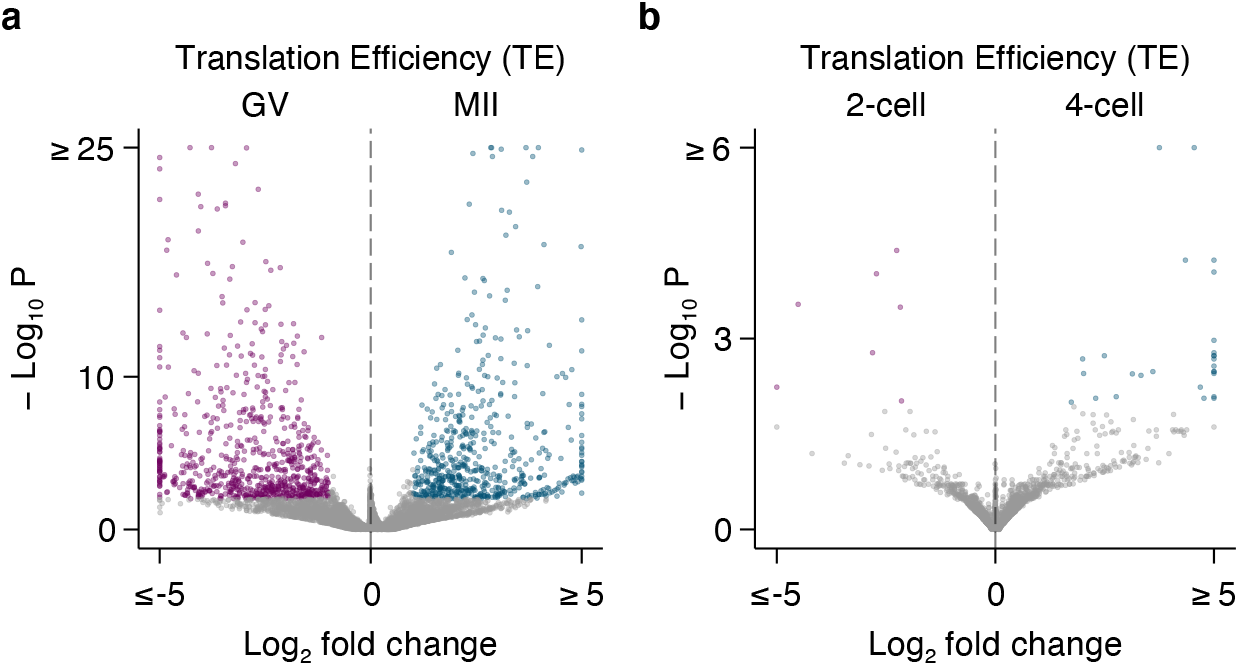
Differential translational efficiency across stages in oocyte and embryonic development. **a, b** Volcano plots depict the statistical significance (y-axis) and log2 fold-change (x-axis) in translation efficiency between the GV- to MII-stage oocytes and between the 2-cell to 4-cell stage embryos. The purple and blue points indicate transcripts with a significant difference in translation efficiency between the compared stages (FDR < 0.01)

